# Drifting Assemblies for Persistent Memory

**DOI:** 10.1101/2020.08.31.276147

**Authors:** Yaroslav Felipe Kalle Kossio, Sven Goedeke, Christian Klos, Raoul-Martin Memmesheimer

## Abstract

Change is ubiquitous in living beings. In particular, the connectome and neural representations can change. Nevertheless behaviors and memories often persist over long times. In a standard model, memories are represented by assemblies of strongly interconnected neurons. For faithful storage these assemblies are assumed to consist of the same neurons over time. Here we propose a contrasting memory model with complete temporal remodeling of assemblies, based on experimentally observed changes of connections and neural representations. The assemblies drift freely as spontaneous synaptic turnover or random activity induce neuron exchange. The gradual exchange allows activity dependent and homeostatic plasticity to conserve the representational structure and keep inputs, outputs and assemblies consistent. This leads to persistent memory. Our findings explain recent experimental results on the temporal evolution of fear memory representations and suggest that memory systems need to be understood in their completeness as individual parts may constantly change.

---

Organisms change over time, on many different levels. This holds in particular for the synapses in neural networks [1]: They change their impact and also appear and vanish. On the one hand, this weight and structural plasticity is activity dependent. Such forms have been argued and directly shown to be crucial for learning [2]. On the other hand, weight changes and turnover with similar strength occur spontaneously, independent of previous spiking activity and in its absence [3, 4, 5, 6]. A similar dichotomy exists for neural representations: They change due to adaptive learning in order to improve task performance, but also spontaneously, often without affecting behavior. The latter has been observed in areas storing long-term memories [7], in sensory areas, for place cells, location and task-selective cells, and in motor areas [8, 9].

Environments change as well. To flexibly adapt, higher animals acquire information and retain it by forming memories in the brain. In a widely used model, a memory is represented by one or several (depending on their definition) neuronal assemblies, i.e. by ensembles of strongly interconnected neurons [10, 11]. For faithful memory storage the ensemble of neurons forming an assembly is assumed to remain the same [12]. Previous theoretical analysis has carefully studied the formation and maintenance of such static neuronal assemblies [13, 14, 15, 16, 17, 18, 19]. In particular, it has been suggested that in presence of noisy, irregular spiking activity [14, 20, 15, 16] and spontaneous synaptic changes [18] assemblies are preserved with the help of activity dependent synaptic plasticity.

Based on the experimentally observed changes of connections and neural representations we develop a contrasting memory model where assemblies are ever and completely changing; they drift. This happens gradually, by successive exchange of individual neurons. The neuron ensembles forming the same assembly at distant time are not directly related, but indirectly via the ensembles forming the assembly at the times in between. Using an analogy of ref. [21] (Supplementary Note), this is comparable to a thread, which consists of many rather short overlapping fibers; the ensembles of fibers in spatially distant parts are not directly related. In our model, the participation of single neurons in the memory representation overlaps, Fig. 1a, like the participation of fibers in the thread. As a consequence, viewed over time the representation looks like a continuous thread, Fig. 1b. While fibers adhere together due to the friction between them, neurons in the assembly adhere due to increased synaptic connection strengths. The inputs and outputs track the course of the “assembly thread” to keep behavior and memory stable, Fig. 1c; they connect at each time to the correct ensemble of neurons that currently forms the required neural representation. Stable input neurons may be located in the sensory periphery, but also within the brain [8], for example in the primary visual cortex and the dentate gyrus, the input area of the hippocampus [22]; motor neurons are candidates for stable output neurons. We will refer to both input and output neurons (Fig. 1c) as periphery neurons and to the assembly forming neurons (Fig. 1a) as interior ones.

**Figure 1:**
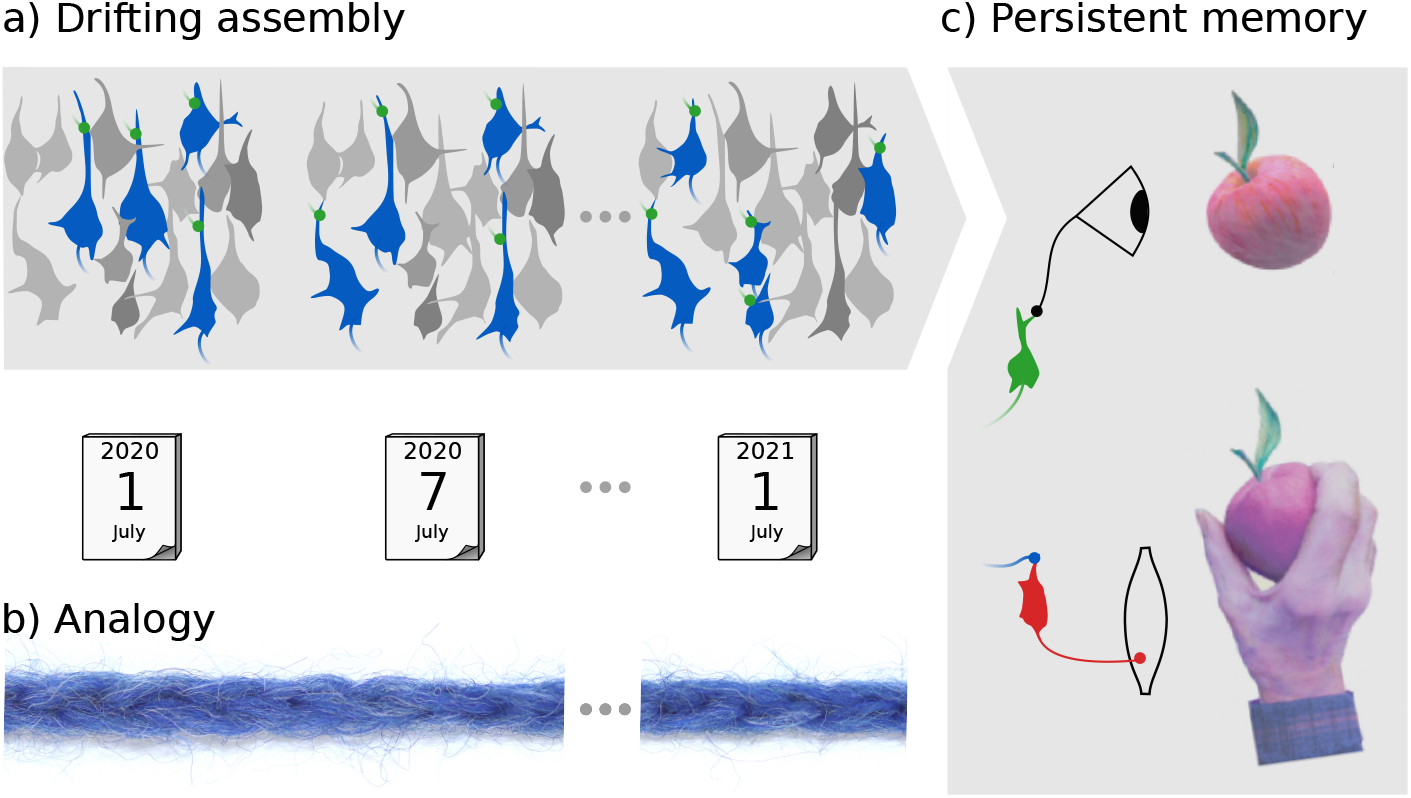
Assembly drift and persistent memory, (a) At two nearby times a similar ensemble of neurons forms the neural representation of, for example, “apple” (compare the blue colored assembly neurons between other, gray colored interior neurons at the first and the second time point). At distant times the representation consists of completely different ensembles (blue colored assembly neurons at the first and the third time point). Due to their gradual change, temporally distant representations are indirectly related via ensembles in the time period between them, (b) Parts of a thread possess the same form of indirect relation: Nearby parts are composed of similar ensembles of fibers, while distant ones consist of different ensembles, which are connected by those in between, (c) The complete change of memory representations still allows for stable behavior. In the figure, a tasty apple is perceived. At different times, this triggers different ensembles that presently form the representation of “apple”, see (a). Assembly activation initiates a reaching movement towards the apple, despite the dissimilarity of the activated neuron ensembles. Memory and behavior are conserved because the gradual change of assembly neurons enables the inputs and outputs to track the neural representation.

We demonstrate the feasibility of our memory model using neural networks at different levels of complexity and with different types of dynamics. Numerical simulations and theoretical analysis reveal the mechanisms leading to the drift of assemblies, the conservation of the overall representational structure and the maintenance of memory. Furthermore, we find that the assembly drift can be directly related to the evolution of fear memory representations uncovered in recent experiments [7] and that they are suitable for computation.

## Results

### Spontaneous remodeling gives rise to drifting assemblies

We show the feasibility of our concept using networks of leaky integrate-and-fire (LIF) model neurons with spontaneous remodeling (Methods). To incorporate the latter, we introduce spontaneous turnover of connectivity, i.e. synapses appear and disappear [3]. The effects of this are easily distinguishable from those of the also present noisy weight changes due to spike timing dependent plasticity (STDP) and noisy spiking. A synapse between two excitatory neurons in our networks has a finite expected lifetime (Methods). Similarly, if the synapse is absent, it has a finite average absence time until it reappears. Excitatory connectivity thus remodels completely.

Figure 2 shows a simulation of a network with 90 interior and 12 periphery neurons. The network is initialized with three assemblies; each has two input and two output neurons. The average life time of a synapse is about half an hour. This is long compared to the membrane and synaptic time constant of individual neurons. Experiments indicate in absence of activity average life times ranging from minutes for immature synapses to two months for mature ones with large weights [3, 5]. We choose shorter lifetimes to reduce the simulation time to a feasible amount and to cover other forms of spontaneous remodeling such as weight remodeling [4]. The presence or absence of the synapse from neuron *j* to *i* is indicated by a one or zero in the entry *c_ij_* of the connectivity matrix of the network. The strength of the synapse is given by the corresponding entry *w_ij_* of the weight matrix. We measure it in terms of the peak excitatory postsynaptic potential (EPSP, units: mV). If not mentioned otherwise, connectivity and weight matrix refers to the matrices between excitatory neurons in the networks, since all other synapses are static. The weight matrix changes when synapses vanish and due to STDP and homeostasis.

**Figure 2:**
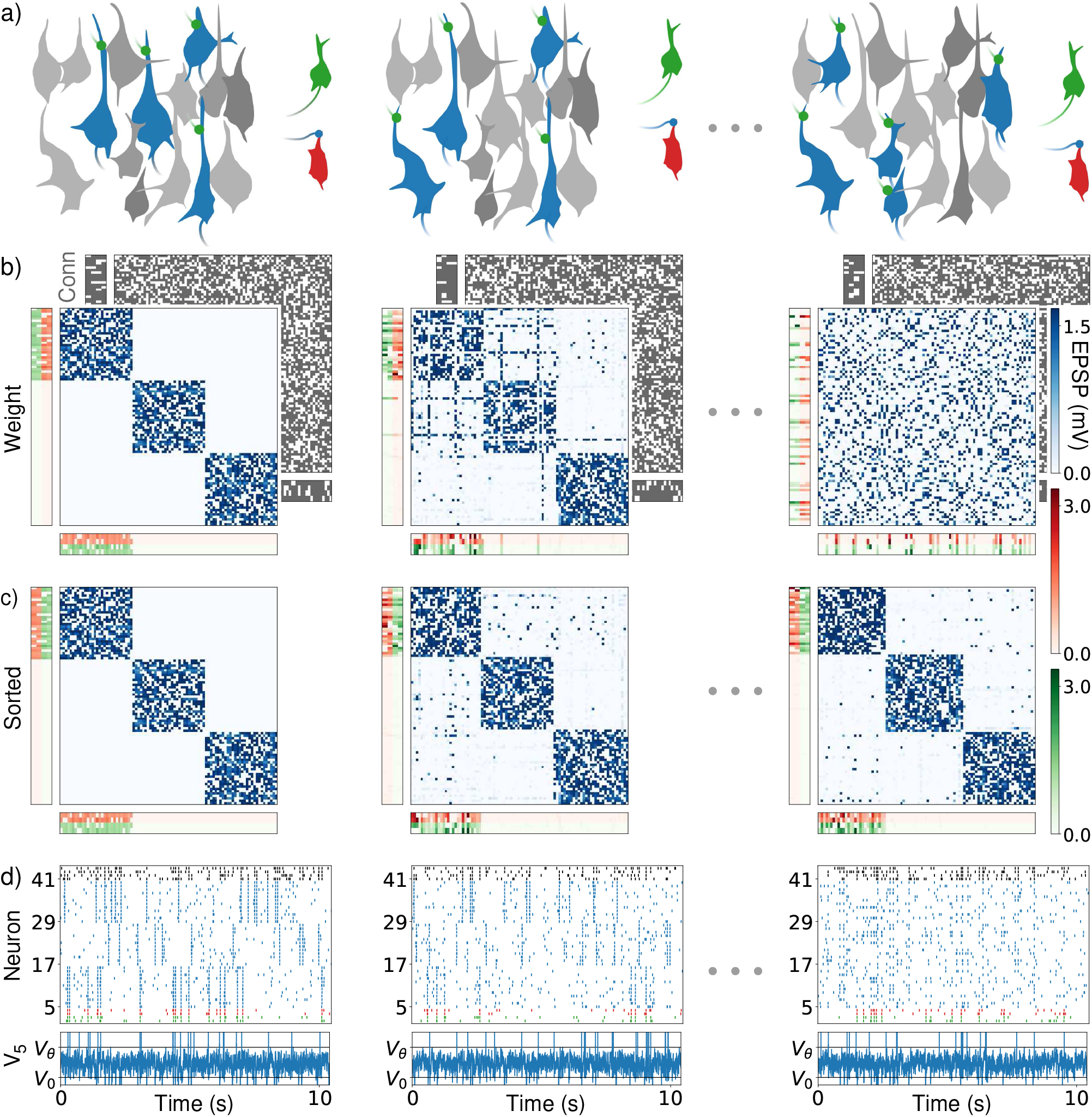
Drifting assemblies driven by spontaneous remodeling, (a) Schematics. While an assembly drifts freely (blue colored assembly neurons) within the interior neurons, its input and output neurons (green and red) follow it by adapting their weights, such that the input-output relation stays conserved, (b) Weights between interior neurons (blue weight matrix), from input and output neurons to the interior neurons (red and green vertical weight matrices) and from the interior to the input and output neurons (red and green horizontal weight matrices). Input (output) weights of neuron i are displayed as ith row (column). Only periphery neurons initially (and thus for all times) attached to assembly 1 are shown for clarity. The underlying connectivity is displayed in the back (gray: present synapses). First column: Network initialization with three assemblies and random connectivity. Second column, after an hour: Remodeling of connectivity has already driven several interior neurons to attach to a new assembly (blue weight matrix, horizontal and vertical “lines” indicating the changed input and output preference). Third column, after 37 hours (h): The assemblies have drifted away, the weight matrix is completely remodeled, (c) Like (b) but with neurons reordered according to assemblies that they belong to, using a clustering algorithm. The three assemblies stay intact and periphery neurons strongly coupled to assembly 1. (d) Spike trains of the input (green) and output (red) neurons of assembly 1, of 12 neurons from each of the ensembles that initially form assembly 1 (5-16), 2 (17-28) and 3 (29-40) and of four inhibitory neurons (black), (d, lower) membrane potential of the first interior neuron. Spikes are marked by vertical lines above threshold V_θ_, reset is to V_0_.

The network simulation shows that the three assemblies drift freely in the network, see Fig. 2b. They spontaneously reactivate, which reinforces them. Background spiking is irregular and asynchronous; the membrane potentials of neurons fluctuate irregularly (Fig. 2d). Reactivations appear dispersed over the entire network at later times, since neurons from all over the network form the three representations. Periphery neurons reactivate together with their assembly, which faithfully binds them to it with strong input and output connections. This keeps memory persistent. Fig. 2c displays the neurons forming assembly 1 with lowest indices, in the upper left corner of the weight matrix. Periphery neurons with strong input from and output to assembly 1 therefore have strong weights at the left part of the horizontal and at the upper part of the vertical weight matrices. We note that drifting assemblies also occur in network models without periphery neurons.

The structural remodeling drives the drift: Without it, assemblies stay invariant after a short time of adaptation to the initial connectivity matrix (Fig. 3, Supplementary Fig. 3). In living animals long-term recordings observed both complete [23] and, more often, incomplete structural remodeling [24]. We will find drifting assemblies also in absence of remodeling, suggesting that they occur in intermediate cases as well, for example when turnover is less random through weight dependence or incomplete [3, 24, 25]. We verified that the drifting assemblies have the “associative memory property” of activating after being partially excited and that they are functional in the sense that after a sufficiently strong stimulation of the input neurons, they activate and stimulate the output neurons (Supplementary Fig. 2).

**Figure 3:**
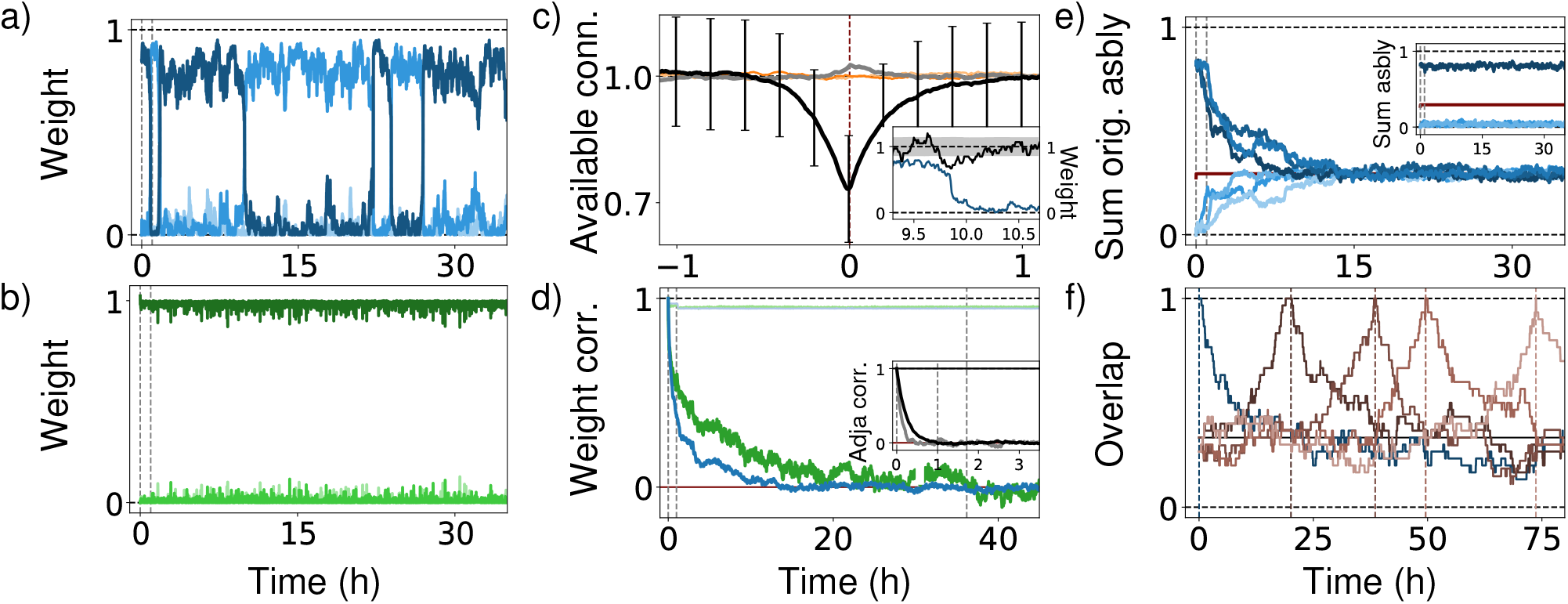
Analysis of drifting assemblies and their periphery neurons, (a) Switching of interior neurons underlies assembly drift. Normalized summed weights between a, neuron and current assemblies 1, 2 and 3 (dark to light blue) show temporary membership and fast transitions. Gray dashed: times used in Fig. 2. (b) Periphery neurons stay attached to their assembly. Display like (a), with greens indicating periphery neuron-assembly weights, (c) Mechanism of switching. Periswitch time histogram shows strong reduction of number of input connections from abandoned assembly near switching (black, error bars: standard deviations of distributions). Input connectivity from entered assembly is slightly increased, (gray). Output connectivity (orange, light orange) has no pronounced trend. Inset: typical switching event. Available input connections (black; black dashed: average; gray: standard, deviation) decay strongly before switching (weight from abandoned, assembly: blue), (d) Complete network weight remodeling. Pearson correlation between initial and, later weights of interior (blue) and periphery-interior synapses (green) converge to chance level (red, 0) Networks with static connectivity undergo initially a, few remodeling steps (light colors). Inset: Same, for the network connectivity, (e) Complete assembly remodeling. Summed, weights within and, between (darker and, lighter blues) the three initial assemblies converge to chance level (1/3). Inset: Maintenance of representational structure. Sum of weights within current assemblies 1 (dark blue) and, between them and, the current other two assemblies (light blues), (f) Continued, assembly drifting. Overlap) of assembly 1 with itself at different reference times: initialization (blue) and, first to fourth complete remodeling time (dark to light brown; times: dashed, verticals).

### Connecting synaptic dynamics and assembly drift

What are the mechanisms underlying the assembly drift and the persistence of memory? Mechanisms of assembly stabilization have been previously described, in static assemblies. They maintain also our assemblies. The highly correlated spiking due to reactivations leads to strengthening of internal and weakening of inter-assembly connections via long term potentiation (LTP) and depression (LTD) [16, 26, 25]. Further, in our model appearing intra-assembly synapses are strengthened and compensate for vanishing ones. Homeostatic plasticity introduces competition, which weakens inter-assembly synapses, as they get potentiated less by STDP [27, 28, 29]. Finally, together with the individual weight limitation homeostasis generates a stabilizing saturation of neurons [28, 16]: Assume that we have only interior neurons in our model. Each neuron then has input synapses and output synapses of a summed strength *w*_sum_ (Methods). An individual synapse has at most weight *w*_max_. A neuron can thus create *w_sum_/*w*_max_* input and output connections of maximal strength. If there are about this many connections with neurons of its own assembly available, the neuron will strengthen them to the limit and hardly generate connections with other neurons. If all its neurons are saturated like this, the assembly is very stable; spontaneously forming assemblies assume approximately this size.

The assembly drift is on the level of single neurons reflected by characteristic dynamics: comparably long times of stable assembly membership, which are interspersed with fast switches between them, Fig. 3a. Analysis of switching events reveals that strong decreases of the total number of available input synapses from its own assembly precede a neuron’s switch as shown by the periswitch time histogram Fig. 3c. Accordingly, networks without structural remodeling show an initial phase of adaptation to the underlying connectivity, where neurons that receive few input synapses from their assembly leave it (Supplementary Fig. 3, Fig. 3d small jump-like decrease in light traces around one hour). The different decay times in Fig. 3d main panel and inset show that synaptic weight plasticity can otherwise compensate large parts of the turnover of connectivity. These observations indicate that in networks with spontaneous remodeling switching neurons are “pushed out” of their assembly: a neuron has a fluctuating number of input connections from its assembly available. If the number becomes too small, homeostasis lets the neuron strengthen its synapses from other assemblies to maintain the desired input level. Supported by weight fluctuations due to noisy spiking, the neuron will then start to spike together with other assemblies and finally perform a fast transition to one of them. This last part of the process will be analyzed below. Neuron switching and thus assembly drift continues over time. Fig. 3 illustrates this by showing that the overlap of realizations of assembly 1 with its initial and with later realizations goes to chance level.

A periphery neuron differs from an interior neuron by higher connectivity, smaller *w*_sum_ and higher *w*_max_ (Methods). It can therefore compensate a higher number of lost input connections from its assembly by increasing the weights of the remaining ones. An interior neuron that newly joins the assembly will start to spike with it and the periphery neuron, such that their connections are strengthened.

### Noisy spiking gives rise to drifting of assemblies

Drifting assemblies also occur in networks without spontaneous synaptic remodeling. To show this, we first use LIF networks with full excitatory-excitatory connectivity and stronger STDP. We find that noisy spiking and the resulting noisy plasticity alone suffice to generate drifting assemblies. This holds for both networks with and without periphery neurons as before. Fig. 4 displays an example for the latter.

**Figure 4:**
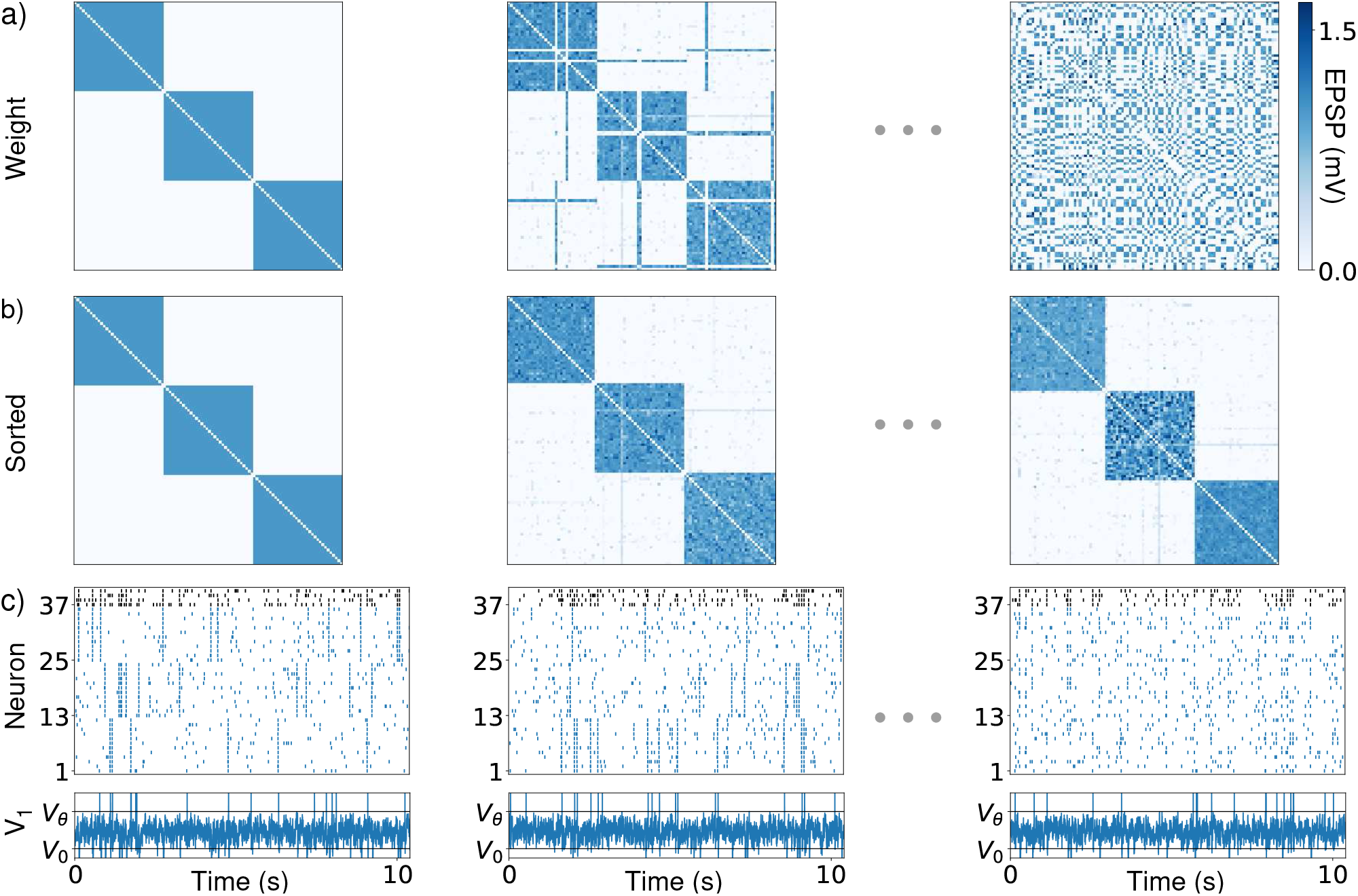
Assembly drift in an LIF neuron network with static connectivity matrix and without periphery neurons, (a) Weights between neurons. First column: Network initialization with three assemblies. Second column, after 15 minutes: first transitions of neurons to a new assembly. Third column, after 12 h: the assemblies have drifted away; the weight matrix has completely remodeled, (b) Like (a) but with neurons reordered according to assemblies that they belong to. The representational structure is conserved over time, (c) Spiketrains of 12 neurons from each of the ensembles that initially form assembly 1 (1-12), 2 (13-2)) and 3 (25-36) and of four inhibitory neurons (black), (c, lower) membrane potential of neuron 1.

To elucidate the mechanism of the underlying neuron transitions we examine the change Δ*w*_1_ of total weight *w*_1_ from assembly 1 to a neuron. We consider only the input weights, since the influence of a single neuron’s output on an assembly is small (cf. also Fig. 3c). Weights are normalized, such that the total input is 1 and therewith *w*_1_ is between 0 and 1. We can now consider all instances where the weight has a certain value, measure the ensuing changes Δ*w*_1_ and compute their average 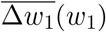 and standard deviation Std(Δ*w*_1_)(*w*_1_) (Methods). If 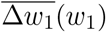 is greater than zero, the input strength from assembly 1 will on average increase: The neuron is “drawn towards” this assembly. This happens for large *w*_1_, Fig. 5b left, i.e. when the neuron is part of assembly 1. *w*_1_ ≈ 0 means that the neuron belongs to assembly 2 or 3 (or, rarely, currently switching between them). In this case *w*_1_ tends to stay away from assembly 1. These dynamics can be visualized by a potential *U*(*w*_1_), where the neuron behaves roughly like an object sliding down its slopes. We see that the average weight change alone lets the neuron stay in the valleys near *w*_1_ ≈ 1 or *w*_1_ ≈ 0, Fig. 5c left. The noise in the plasticity, quantified by Std(Δ*w*_1_)(*w*_1_) in Fig. 5d left, is thus crucial for transitions, like in noise-activated transitions between meta-stable states [30]. Interestingly, already the presence of strong noise in the transition zone lets the neuron choose one of the assemblies (Supplementary Note) and thereby creates meta-stable states (noise-induced multistability) [31].

**Figure 5:**
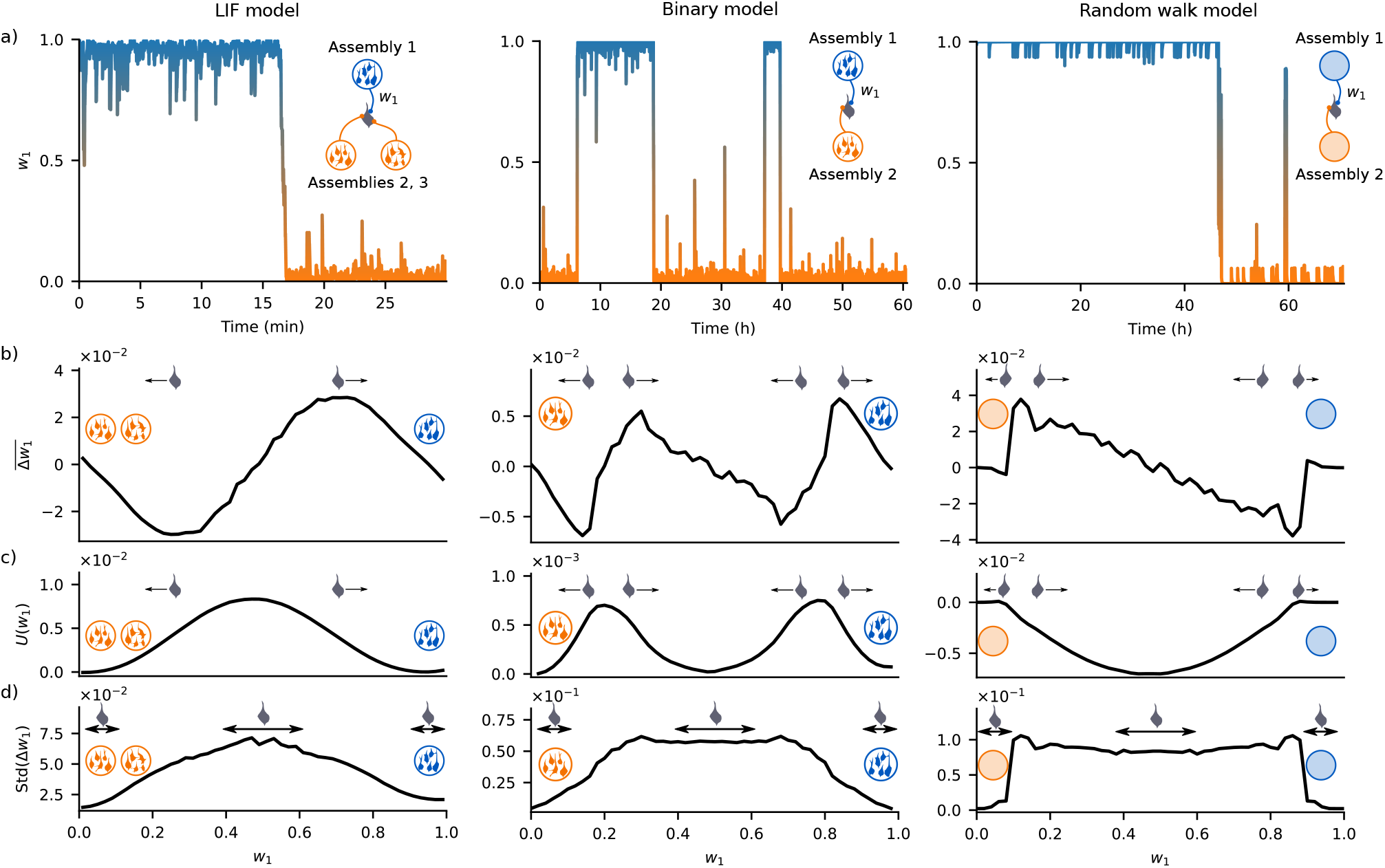
Mechanisms of neuron transitions between assemblies due to noisy activity, STDP and homeostasis. The figure shows similar but in details different mechanisms for the LIF (left column), binary (middle) and random walk (right) models, (a) Longer-term membership of a, neuron in assemblies and fast transition between them. Large total input weight w_1_ from assembly 1 to the neuron (blue shading, in contrast to orange) reflects membership in this assembly, (b) Average change 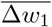 of the total input weight w_1_ from assembly 1, as a function of the current weight value, (c) Effective potential U(w_1_) for the average weight changes. The average weight changes are roughly similar to the displacement of an object sliding down the landscape given by U(w_1_). The two valleys near 0 and 1 are meta-stable states corresponding to different assembly memberships, (d) Standard deviation Std(Δw_1_) of the change of the total input weight w_1_ from assembly 1, as a function of the current weight value.

The above insights suggest to add the following detail to the explanation of neuron switching in our networks with spontaneous remodeling, Fig. 2. There, the fluctuating sparse connectivity introduces a fluctuating bound on the total synaptic input weight that a neuron may receive from its current assembly. Downward fluctuations of this bound can bring an interior neuron so close to the edge of its current potential valley that the otherwise insufficient plasticity fluctuations are able to induce a transition to another assembly.

To enable long-term simulations for a comparison with experiments, we consider a binary model with weight-dependent covariance learning (Methods), which also shows assembly drift without underlying spontaneous remodeling. Its remodeling dynamics resemble those of the LIF networks (Supplementary Fig. 4). We study the underlying switching mechanism in a network with two assemblies for simplicity.

It turns out to be slightly different than in the LIF model, as there is an additional valley in the middle of transition, Fig. 5 middle. The strong noise in this regions prevents the neuron from getting trapped there. The switching dynamics of the binary system can be reduced to an effective one-dimensional random walk, Fig. 5 middle (Methods). This reduced model shows that the additional potential valley (Fig. 5b right) occurs because the neuron spikes in the transition zone with both assemblies with equal probability and homeostatic normalization favors the assembly with smaller weight, preventing the emergence of selectivity. The spiking with both assemblies also induces the high noise in the weight changes. Eventually, one assembly, by chance, induces more changes than the other; the neuron reaches that assembly’s potential and noise valley and settles down.

If present, periphery neurons in the networks without spontaneous remodeling have synapses that are less plastic compared to those between interior neurons. Therefore they react less sensitively to fluctuations in the spiking activity and do not leave the potential valleys of their assemblies. In the networks with spontaneous remodeling the downward fluctuations of the input weight bound stay too far from the potential valley edge for transitions to happen.

### A model for experimentally found evolution of fear memory representations

The evolution of a long-term memory representation was recently experimentally investigated in the prelimbic cortex [7], a structure known to be crucial for context dependent aversive memory [32]. In the experiment mice underwent context dependent fear conditioning with an auditory stimulus. This was followed by two retrievals, the first after 1, 7 or 14 days, the second after 28 days. Neurons active during fear conditioning or the first retrieval were detected by using a genetic marker, TRAP2, and neurons active during the second retrieval by utilizing Fos expression. The numbers of neurons forming the memory representation were approximately the same in all sessions, as well as the behavioral response (freezing) during retrievals. In contrast, the subset of neurons labeled by both markers and thus the overlap of the neuron ensembles forming the memory representation, increased with decreasing time between the two labeling sessions, Fig. 6b, left.

**Figure 6:**
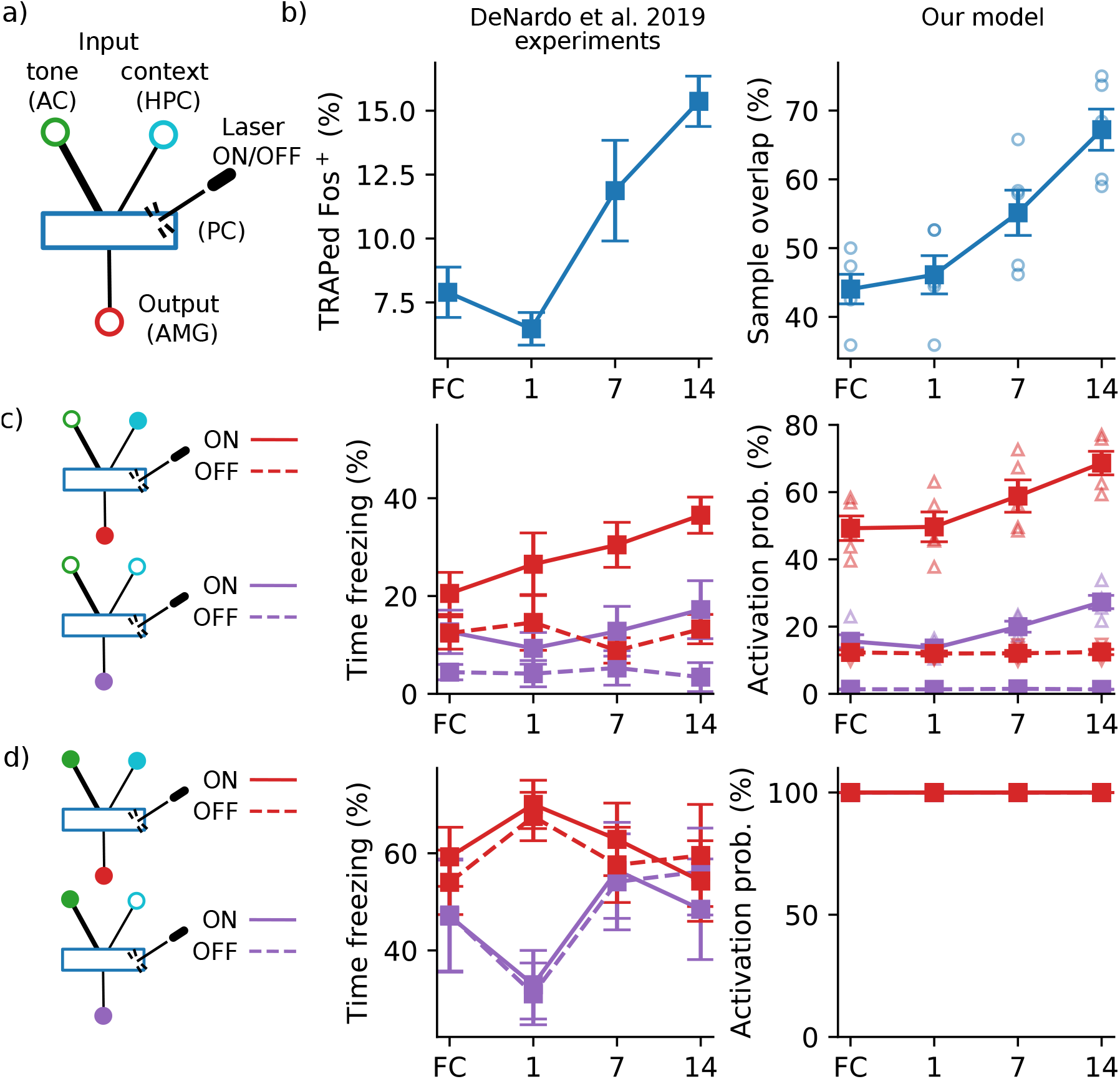
Evolution of fear memory representation observed in ref. [7] and drifting assembly model, (a) Model for the fear memory circuit with context input signaling conditioned cage from hippocampus (HPC), auditory stimulus input signaling conditioned tone from auditory cortex (AC), output to amygdala (AMG), and connecting drifting assembly in prelimbic cortex (PL). In some experiments a set of neurons in PL is photostimulated. Thicker lines visualize stronger connections, (b, left) Experimental data that are proportional to the overlap of the memory representation at the second retrieval (day 28, Fos^+^ neurons) with that directly after conditioning (FC) or during first retrieval, on day 1, 7 or í) (TRAPed neurons). The panel displays the fraction of Fos ^+^ neurons that were also TRAPed, in percent, (b, right) Overlap) of the memory assembly in our model on day 28 with that at the beginning of the trial (FC), and on days 1, 7 and 1) (circles; squares connected by solid lines: mean), (c) Fear expression tested on day 28 in the conditioning context (red, context input on) and on day 29 in an alternate context (purple, context input off). Laser activating a sample of the memory representation labeled (TRAPed) on different days as in (b) was either on (upward triangles; squares, solid: mean) or off (downward triangles; squares, dashed: mean). Left: experiment; right: our model, (d) Same as (c) but in presence of conditioned auditory stimulus (tone input on). Red continuous lines overlay dashed and purple ones on the right. Graphs show mean ± s.e.m and data points.

**Figure 7.**
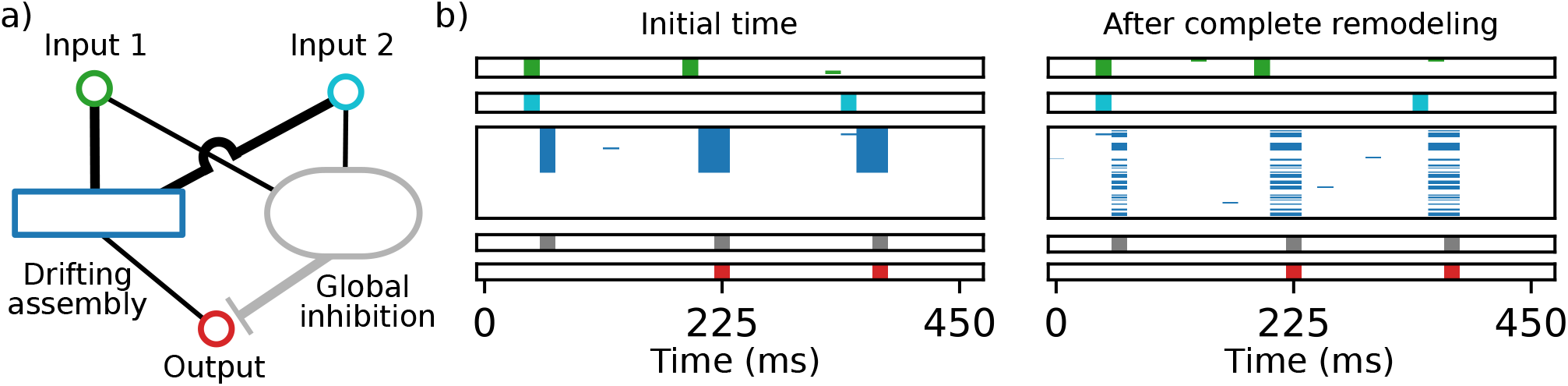
Drifting XOR gate, (a) Schematics of the network setup. Two groups of input neurons (green and cyan) are connected via a drifting assembly (blue) to output neurons (red). An inhibitory population (gray) inhibits all neurons. Thicker lines illustrate stronger connection. Only projections and neuron populations that are directly relevant for the XOR computation are shown for clarity, (b) Functionality of the drifting XOR gate at two distant times. Activity of input 1 (green), input 2 (cyan), hidden layer (blue), inhibitory (gray) and output (red) neurons is displayed by horizontal lines in different bars (from top to bottom). The interiorneurons are indexed such that initially (left) the XOR assembly activity fills the upper part of the third bar. Later, due to complete remodeling of the assembly, its activity is distributed all over the bar (right). Functionality is nevertheless conserved: the output neurons become active if exactly one of the input neuron groups is active.

We conjectured that the experimental findings can be explained if we assume that a drifting assembly forms the memory representation. To model the system, we combine an assembly in prelimbic cortex with “context” input neurons in the hippocampus and auditory stimulus “tone” input neurons in the auditory cortex. These neurons signal that the animal is in the cage where fear conditioning had taken place and that the conditioned tone is played. The assembly’s output neurons are in the amygdala [33], Fig. 6a (Methods). We take the probability of their activation as indicator of the animal’s freezing time. In an animal, their activation is linked to the freezing response via internal amygdala processing and downstream areas, which we assume to limit the freezing response to the observed 60% of the time and to ensure that it occurs in continuous, longer stretches [32].

The model replicates the main experimental findings: The overlap between the neural representations of memory decreases for longer periods between sessions Fig.6b right, since more assembly neurons switch. The assembly sizes are on average unchanged, as the assemblies only drift. Finally, the behavioral response is conserved, because the input and output neurons faithfully follow the drift.

The model also accounts for more details of the experimental results. The conditioned tone alone induced more freezing than the conditioned context alone. The model replicates this (Fig. 6c,d right, dashed lines), since the tone in auditory cortex is represented by more input neurons than the context in hippocampus. The experiments included photostimulation of neurons that were labeled during fear conditioning or the first of the two retrieval sessions. In absence of the conditioned tone, this affected behavior the stronger the smaller the period between retrieval and stimulation was, Fig. 6c,d left, solid lines. Our model captures this result, since the more time has passed since labeling, the more of the labeled neurons have transitioned away from the assembly. The laser thus stimulates a smaller part of the current assembly and activates it more rarely, which is reflected in the output, Fig. 6c,d right, solid lines. In both experiment and model, the conditioned tone input has a strong impact on behavior and results in a similar level of fear expression with and without photostimulation, in the conditioned as well as in the alternative context, Fig. 6d. We finally note that there is freezing in absence of any stimulus, Fig. 6c left. Our model replicates this since the assembly spontaneously reactivates, which also activates the output neurons.

Our results suggest that the changes in ref. [7] occur due to gradual replacement of neurons in strongly connected ensembles, driven by spontaneous and noise-induced plasticity, which can possibly be modulated. The model predicts a continued decay of overlap to the chance level of the coding density, i.e. the fraction of neurons active during assembly activation (see Fig. 3f where it is 1/3). Specifically, this indicates that the fraction of Fos stained neurons of previously TRAPed ones (Fig. 2e of [7]) will go down to 8% (coding density, Supplementary Fig. 4d in [7]) in an a few weeks longer experiment. Remodeling will continue also thereafter, Fig. 3f. Finally, the model predicts that synaptic weights completely remodel, Fig. 3d. Drifting assemblies may also explain the representational change observed in [34], with partial stabilization over time resulting from reduced novelty and thus plasticity or excitability. The implementation of assemblies will likely differ, as recurrent excitation is rare in the considered brain area; the experimental predictions are nevertheless similar.

### Computation with drifting assemblies

Drifting assemblies enable non-trivial computations. We show this by example of a XOR gate, a classical example of a problem that is not linearly separable. Such problems cannot be solved by a simple single layer neural network (perceptron), which sums its weighted inputs and applies a threshold function. XOR has two inputs and an output. If exactly one of the inputs is active, the output gets active. Figure 6a shows the XOR computation realized by a circuit of binary neurons with drifting assemblies in the hidden layer between inputs and outputs. The circuit works as follows: If a single of the two groups of XOR input neurons is activated, the XOR assembly gets active in the next step. This activates in the third step all XOR output neurons. Simultaneously it activates an inhibitory population, which suppresses the activity in the network in the step thereafter. If both sets of XOR inputs activate together, in the second step the XOR assembly activates as well. The input is, however, strong enough to also directly activate the inhibitory population. Consequently, activity in the network is suppressed in the third step. In particular the output neurons remain silent. Due to the spontaneous activity in the circuit, the XOR assembly drifts and the periphery neurons follow it, which preserves the XOR input-output functionality over time, Figure 6b.

Spontaneous background activity leads occasionally to the activation of assemblies and thereby also of the output neurons. These spontaneous events may occur only during certain brain states, such that they have no influence on behavior. Furthermore, real circuits may be composed of many assemblies, such that spontaneous reactivation of one is not enough to trigger reactivation of the entire circuit. Finally, meaningful inputs will usually occur for longer periods, such that also the output stays repeatedly active over a longer period. A short, spontaneous output activation may thus be a signal that is simply ignored. Alternatively, the spontaneous reactivation may account for occasional erroneous behavior, such as spontaneous freezing.

### Drifting assemblies in networks without spontaneous assembly reactivation

In the LIF and binary neural network models considered so far, the assemblies undergo spontaneous reactivations: events during which practically all neurons in the assembly are active. Such reactivations have been observed in various cortical areas, from visual and auditory cortex to the hippocampus [35]. To address their necessity for drifting assemblies, we use networks of linear Poisson (or “Hawkes”) model neurons [36, 37, 16, 38]. The synapses in our Poisson networks change according to a plasticity rule that does not require pre- and postsynaptic spiking, but already generates potentiation if one neuron spikes and its partner is more than on average excited [39, 40] (Methods). Furthermore, the synapses spontaneously remodel. This drives assembly drift as in Fig. 2, Supplementary Fig. 6.

Assembly structure and drift are like those observed in LIF and binary network models. The pairwise correlations in the Poisson model are, however, much smaller (Fig. 8 columns one and two). They are also much smaller than in the brain [41]. (The large correlations in the LIF and in binary networks are mainly due to the frequent reactivation of individual assemblies, Fig. 8a,b third column, Supplementary Fig. 7.)

**Figure 8:**
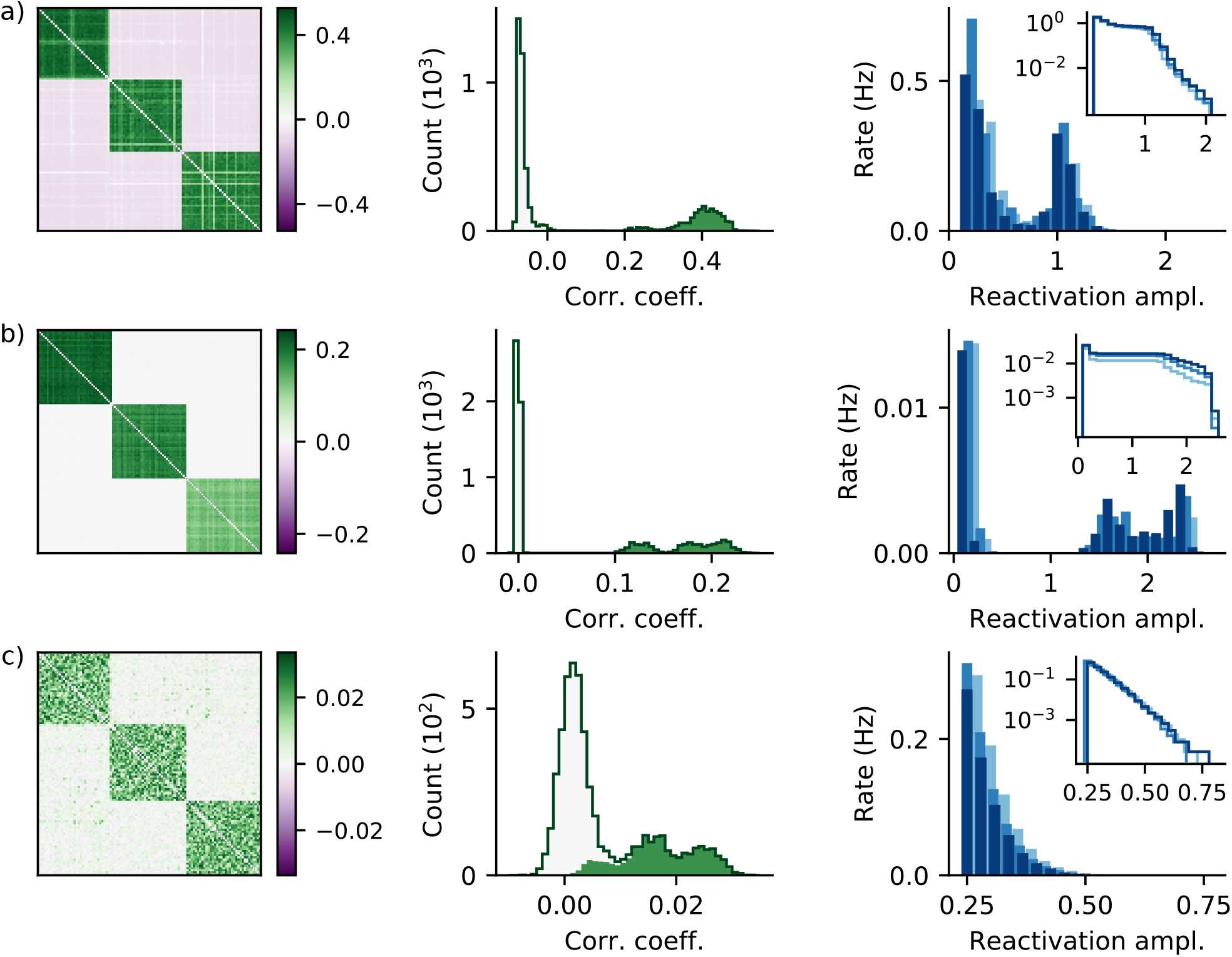
Spike correlations and assembly reactivation in our network models, (a) LIF network of Fig. 2, right column, (b) Binary network with three assemblies (see Supplementary Fig. 4). (c) Poisson network with three assemblies (see Supplementary Fig. 6). All networks have frozen connectivity and weights, which are obtained by fixing plastic networks after the first complete remodeling. First two columns: Correlations between Poisson neurons (c) are much smaller than those in the other two networks (a,b). First column: Matrices of measured spike count correlation coefficients with neurons sorted according to their assembly membership. Diagonal entries (equal to one) are blanked in white. Second column: Histograms of correlation coefficients in first column. Solid line gives the full histogram over all neuron pairs, green shading indicates the contribution from intra-assembly neuron pairs, light shading the remaining contribution from pairs of neurons belonging to different assemblies. Third column: Assembly reactivation is absent in Poisson networks. Main panels show how often different maxima of the number of spikes in a moving time window were detected within an assembly (see Methods). Occurrence rate is plotted against number of spikes divided by the assembly size (different blues for the three assemblies). Events of spontaneous assembly reactivation are reflected by a second peak, separate from the background continuum near zero. Insets show the complementary cumulative distributions with logarithmic rate scale.

Ignition-like reactivation like in the LIF (Supplementary Fig. 2) and binary model can in principle not occur in networks of linear Poisson neurons, since each spike has the same impact. In contrast the dynamics consist of overlapping “avalanches”, transient sequences of spikes evoked by a spontaneous one [38]. For small total synaptic weights of neurons, these are short-lived and small. An avalanche usually activates a part of a single assembly, since the interconnections there are still comparably large. Avalanches involving entire assemblies do not occur on relevant time scales (Fig. 8c right). We observe that the occurring partial activation already suffices to sustain a drifting assembly: Its inactive neurons also have an increased level of excitation, due to the received spikes. The resulting potentiation between inactive and active neurons adds to that between the active ones. Both contributions together keep the assemblies intact and let neurons switch in conjunction with synaptic turnover.

## Discussion

Our work proposes a model for memory that is based on drifting assemblies. The drift results from fast transitions of neurons between assemblies, which are generated by spontaneous synaptic remodeling and irregular spiking. The represented memories nevertheless persist, because STDP and homeostasis are able to keep the representational structure intact and inputs, outputs and the assemblies consistent. This happens without requiring teaching signals or behavioral feedback, even though the network connectivity, the weight matrices and the ensembles of neurons forming the assemblies change completely. The drifting assemblies are suitable for computations and they explain recent experimental findings on changes of memory representations.

Several general suggestions have been raised on how neural network functionality may be maintained despite spontaneous synaptic remodeling and unstable neural representations: First, the changes might have no impact on the relevant part of network dynamics because they are too weak or because they are eliminated by downstream attractor dynamics [1, 42, 8]. Secondly, spontaneous network remodeling may generate transitions between redundant networks that are equally well suited for the required tasks [43, 1, 42]. Thirdly, sensory feedback or feedback from other brain areas may lead to the correction of remodeling that is not in redundant directions [42, 9]. Previous computational studies addressing the impact of intrinsic synaptic fluctuations focused often on the question how neural representations can nevertheless be stable. They proposed that a preserved core structure keeps neuronal activity stable [44, 8, 45, 46], that the networks are retrained to counteract degradation [43, 47, 48] or that the ensembles of neurons storing memories are kept invariant via reactivations and unsupervised learning [49, 18, 25]. A few computational studies addressed how representations that change may emerge and partially also how the change’s impact may be attenuated. Ref. [43] finds that the synaptic remodeling lets the preferred directions of hidden layer model neurons fluctuate similar to preferred directions in some recordings of motor cortex neurons. Ref. [50] shows that modest retraining of linear readouts by supervised learning allows to detect location, speed and head direction from changing representations in the posterior parietal cortex. Ref. [51] found in a model for developing zebrafish networks spontaneously emerging and changing assemblies, which break down and merge and thus seem unsuitable for persistent representation. Ref. [46] shows that if a fluctuating part is added to otherwise stable synaptic weights storing a sequence, individual neurons partially and temporarily change their dynamics, but the original sequence can still be detected.

In our networks there is no preserved core of network structure or memory representation. Only the full circuit consisting of inputs, assemblies and outputs preserves memory and behavior. Interior neurons switch from coding for one task to coding for another, only periphery neurons have stable codes. The inputs and the readouts therefore have to constantly compensate for remodeling, by unsupervised plasticity combining STDP and homeostasis. This requires no external feedback and no error signal exchange within the brain, as collective activity tells the neurons where to attach. Such unsupervised compensation may occur for other types of drifting representations as well. We note that a corrective feedback from other brain areas would require a stable anchoring in an invariant representation, otherwise there will be an error in the error signal, thus co-evolution of errors and finally erroneous behavior. On the level of synapses there is constant change in both periphery and interior neurons in our model; no synapse is persistent. This predicts that in memory areas weight remodeling will turn out to be complete [6].

The change of networks and representations through assembly drift is functionally per se neutral. Modification and acquisition of memories will happen on top. The drift might nevertheless help the storage of memories and contribute to solving the stability-plasticity dilemma: New memories can be imprinted in the same highly plastic (sub-)region (cf. also the two-stage model for memory [10]). With time their representations drift away together with their input and output connections, giving way to further ones.

The developed models combine established model neurons [36, 13, 11], simple STDP or correlation learning [11, 16, 26] and homeostasis [28, 16, 51, 29]. Robustness is indicated by the occurrence of drifting assemblies in different neuron models, the simple mechanisms underlying the drift and the broad range of correlation strengths that can sustain drifting assemblies; the range includes the correlations found in biological neural networks [41]. Our models can generate very different drifting times from hours to years, which depend on underlying parameters like noise strength and synaptic lifetime. Furthermore, the drifting of assemblies is consistent with the requirement of executing non-trivial computations with them. Finally, assuming that drifting assemblies form the memory representations in the prelimbic cortex of mice explains the main and several detailed experimental findings in ref. [7]. We therefore propose that assembly drift as predicted by our models occurs in biological neural networks. Our findings suggest a specific connection between spontaneous synaptic remodeling and the change of neural representations: remodeling pushes individual neurons out of their assembly and lets them switch to new ones. We find that noisy spiking and the resulting noisy plasticity alone can also cause the transitions of neurons between assemblies. This is due to noise driven switching between metastable states and inhomogeneous noise that prevents neurons to linger between assemblies. Since in biological neural networks spontaneous remodeling is strong [3, 4, 6] and spike noise high [11, 41], we suggest that both together drive the drifting of assemblies in biological neural networks. The strength of input connections thereby decides whether an interior neuron continues to stay with its assembly, suggesting that input neurons rely on feedback from their assembly to be able to follow it. Our model thus provides an additional explanation for the abundance of feedback connections throughout cortical areas, including early sensory ones.

Our results make specific and general predictions for memory systems. In particular, they predict a continued drift, reduction of overlap to chance level and gradual but complete remodeling of synaptic weights, as well as the mechanisms underlying the drift and ways to interfere with it. On a more general level, our results indicate that impairments of synaptic plasticity do not only lead to problems in acquiring new memories but also to forgetting, either because assemblies break down or because the improperly compensating inputs and outputs disconnect from their drifting assemblies. Changes in synaptic plasticity as observed in dementia may thus directly lead to the symptomatic forgetting. Further, unsupervised compensation of drifting representations could be a common theme in the brain. Finally, our work suggests that Heraclitus of Ephesus’ 2500 year old idea is also true for memory representation, namely that: 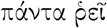; everything flows or, in other words: drifts.

## Methods

### Models and approaches

To demonstrate the feasibility of drifting assemblies we consider networks of LIF model neurons. These incorporate important features of biological neurons such as interaction with spikes in continuous time, reset after spiking and integration of input. Simultaneously, they describe neuronal dynamics with comparably few variables and parameters, which facilitates the long-term simulations required for our study and the model setup. The long-term simulations also require that we consider networks of medium size, of the order of a hundred neurons. For more specific questions, for example to elucidate the details of dynamical mechanisms, we use further simplified models of Poisson spiking [36, 37, 16, 38] or binary neurons [13, 28, 51, 29]. Since the memories are in our model stored in the synapses between excitatory neurons, only these are plastic. Previous theoretical work has shown that learning [15, 20] or spontaneous emergence [16, 19] of static assemblies as well as their maintenance can be enabled by a combination of STDP and homeostatic plasticity. A recent study investigated STDP in the recurrent excitatory synapses of the hippocampal region CA3 [26], i.e. of a region that is assumed to serve as an associative memory network and store assemblies [10]. It found an STDP learning window with LTP for pre- and post-synaptic spikes, irrespective of their ordering and the stronger the closer they are. We thus choose in our models an STDP rule with a symmetric learning window with centrally peaked LTP (Supplementary Fig. 1). LTP is compensated by LTD for temporally distant spikes, see also [16]. In networks of binary neurons we reduce the STDP to a standard Hebbian covariance rule [11, 51] and augment it by a dependence of weight changes on previous weight. Finally, in our networks of Poisson neurons we use an STDP rule that is similar to voltage- or calcium-based rules [39, 40]. In all models the synapses further undergo homeostasis. Experiments indicate that the total input [52, 28, 53] to a neuron may be conserved. Further, there is evidence for output normalization [54]. Both input and output normalization may be realized by competition for synaptic resources [28]. Following [28, 16, 51, 29], we thus introduce normalizations of the model neuron’s input and output weights. Finally, we incorporate spontaneous remodeling in some of the models since we expect it to be a possible source of assembly drift.

### LIF networks

We use a current-based LIF neuron model. The membrane potential *V_i_*(*t*) of neuron *i* obeys

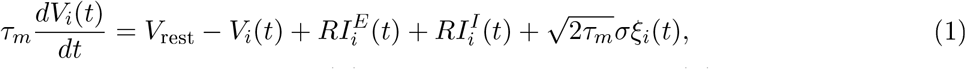

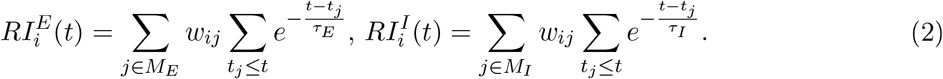

Here, *τ_m_* is the membrane time constant, *V*_rest_ the resting membrane potential, *R* the input resistance, 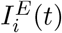 and 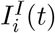 are the total currents generated by the populations of excitatory and inhibitory synapses and *ξ_i_*(*t*) is standard Gaussian white noise. The parameter *σ* equals the standard deviation of the membrane potential’s stationary distribution in absence of a threshold and synaptic input currents from the other modeled neurons; the membrane potential then follows an Ornstein-Uhlenbeck process. When the voltage exceeds a spike threshold *V_θ_* = 20mV, a spike is generated and the neuron is reset to Vo = 0mV, where it stays for a refractory period *τ*_ref_. *V*_rest_ is halfway between threshold and reset. A generated spike travels to postsynaptic neurons, where it generates changes in the synaptic currents. A spike of an excitatory neuron j evokes in the input 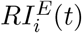 a jump-like increase of height *w_ij_*, which thereafter decays exponentially with time constant *τ_E_*. The decay time constant of inhibitory input currents is *τ_I_*. *t_j_* are the spike times of neuron *j*, *M_E_* is the set of all *N_E_* excitatory and *M_I_* that of all *N_I_* inhibitory neurons.

At each excitatory spike, the weights are updated according to a pair-based spike rule with symmetric STDP window (Supplementary Fig. 1). Each side of the window is the difference of two exponentials with decay time constants τ_LTP_ = 20ms and τ_LTP_ = 40ms. Peak LTP is 1.25mV (3.75mV) at 0ms, peak LTD is −0.42mV (—1.25mV) at ±44ms for networks with (without) periphery neurons and turnover. Homeostatic plasticity normalizes the total excitatory input and output strength of an interior neuron i to ∑_*j*∈*M_E_*_ *W_ij_* = Σ_*j*∈*M_E_*_ *w_ji_* = *w*_sum_; for a periphery neuron the total input and output weight strength is *w*_sum,peri_. The normalization is approximated by normalizing after each excitatory spike the columns and the rows of the weight matrix between the excitatory neurons. Excitatory couplings between interior neurons are bounded by 0mV and *w*_max_, for couplings from or to periphery neurons the upper bound is *w*_max,pc_. This bound is enforced by clipping weights before and after homeostatic normalization. All possible synapses from excitatory to inhibitory neurons are present, they are non-plastic and have identical weights. The same holds for synapses from inhibitory to excitatory and inhibitory neurons. Periphery neurons are not interconnected. There are no self-couplings.

### Assemblies

Memory representations in our model are drifting assemblies: We propose that they gradually change over time by replacing their neurons. An assembly nevertheless maintains its identity to allow for stable memory, Fig. 1. In particular, it does not vanish, split or merge with others. Bridging from memory to behavior, the assemblies have input and output neurons. If these are present, a particular set of them needs to follow an assembly and thereby allows us to identify it over time, in addition to the identification by indirect resemblance. Persistent memory manifests itself as a conserved behavioral input-output relation: the input neurons that activate the different ensembles forming an assembly are not exchanged over time as well as the output neurons activated by them. For simplicity, we assume that memories and the related behaviors are unchanged over time; the subject does not for example learn more about apples to modify behavior. We further assume that assemblies do not share neurons and each input and output neuron is specific to one representation. Networks are initialized by setting existing weights between neurons within an assembly and between an assembly and its periphery neurons to 1mV and all others to 0mV. Thereafter outputs and inputs of each neuron are normalized to *w*_sum_ or *w*_sum,peri_ and individual connections are clipped to their allowed ranges.

### Connectivity remodeling

A synapse between two excitatory neurons *j* (presynaptic neuron) and *i* (postsynaptic neuron) in our networks with spontaneously remodeling has a finite expected lifetime *L*. It vanishes with rate 1/*L*, i.e. in a simulation step of duration *Δt* with probability Δ*t/L*, independent of activity. Similarly, if the synapse is absent, it has an average absence time *A*; it appears in a simulation step with probability Δ*t/A*. Thus, on average the synapse is present a fraction *L/*(*L* + *A*) of the time; the probability that it is present at a certain time point is *p* = *L*/(*L* + *A*), the density of synapses of a certain type. The spontaneous remodeling in our networks switches entries of the connectivity matrix between 1 and 0. When a synapse vanishes, its weight becomes 0mV. Newly appearing synapses have weight 0mV as well [5].

### Simulation of LIF networks and analysis

The network in Fig. 2 consists of *N_E_* = 102 excitatory and *N_I_* = 20 inhibitory neurons. *N*_int_ = 90 of the excitatory neurons are interior neurons. These are initiated such that there are three assemblies of *N*_asbly_(0) = 30 interior neurons. Each assembly has 4 periphery neurons. The first two periphery neurons are designated input, the second two output neurons. The connection density is *p*_int_ = 0.6 between interior neurons and *p*_peri_ = 0.8 between interior and periphery neurons. The density of present synapses is high to compensate small network size. The corresponding synaptic life and absence times are for synapses between interior neurons *L*_int_ = 2000s and *A*_int_ = 1333.3s and for those between interior and periphery neurons *L*_peri_ = 2000s and *A*_peri_ = 500s. Total simulation time is 100 hours. Four periphery neuron weights exceed the range of the colorbar in Fig. 2; the largest weight would evoke a 3.9mV high EPSP. We did five alike simulations with different random realizations of networks and noise to check that the representational structure is conserved over time, i.e. that assemblies continuously drift and their periphery neurons faithfully follow them. Clustering is throughout the article obtained with the Louvain clustering algorithm [55] as implemented in [56].

Fig. 3a shows the sum of the weights between interior neuron 1 (indexed 5 in Fig. 2d) and assemblies 1,2 and 3, normalized by 2*w*_sum_ (total input plus total output weight). Dashed black horizontals display maximal and minimal sums of weights, (b) similarly shows the sum of the weights between input neuron 1 and assemblies 1,2 and 3 normalized by 2*w*_sum,peri_. To obtain the black curve in the periswitch time histogram (c) we collect the numbers of available input connections from the abandoned assembly and its periphery neurons to the switching neuron, for all switching events in the simulation of Fig. 2. We then normalize them by their expected number using the current number of assembly neurons and the connection probabilities *p*_int_ and *p*_peri_. Thereafter, the collected pieces are centered at switching time, which is set to zero (red dashed vertical). The numbers of available input connections at a time point are then averaged and their standard deviation is computed. The other curves are computed likewise. The inset shows the third switching of assembly neuron 1, away from assembly 1, at about ten hours simulated time (red dashed vertical, cf. panel a). (d) The Pearson correlation between weight matrices at times 0 and t is computed as 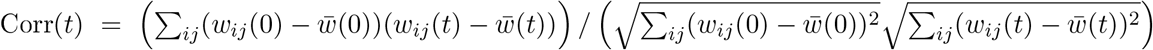, where 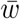 is the average entry size. The expression for connectivity matrices is analogous. The maximal correlation is 1 (dashed black horizontal line) (e) Displayed is the sum of the weights between the neurons originally forming the first, second, third assembly (dark shadings of blue), normalized by their maximal sum *N*_asbly_(0)*w*_sum_. Further displayed is the sum of weights between the neurons originally forming the first and second, first and third, second and third assembly (light shadings of blue), normalized by their maximal sum 2*N*_asbly_(0)*w*_sum_. The inset shows the analogous quantities for the current assemblies at each time, normalized using the current assembly sizes. Chance level at a time is computed by summing all weights between interior neurons and normalizing by *N*_int_ *w*_sum_. Dashed lines indicate maximal and minimal sum. Panel (f) shows the overlap of the realizations of assembly 1 with previous and future reference ensembles. We compute the overlap as the number of neurons that an ensemble of neurons shares with its reference ensemble, normalized by the size of the reference ensemble. The overlap is thus bounded by 0 and 1 (dashed black horizontal lines). The first reference in Fig. 3f is the ensemble initially forming assembly 1. We observe complete remodeling (blue trace), i.e. the overlap with the reference decreases to chance level. Setting the realization of assembly 1 at the time when the overlap reaches there (first complete remodeling time) as second reference ensemble reveals another complete remodeling thereafter (brown trace). It follows a third remodeling (lighter brown trace) etc. Due to the continuous spontaneous remodeling over the entire course of the simulation, also the neurons’ switching continues and the assemblies remodel again and again. We use as criterion for complete network remodeling that for all assemblies the overlap with their original realization has reached chance level at least once.

In addition to stronger STDP, the network in Fig. 4 differs from the network in Fig. 2 in the following aspects: all of its excitatory neurons are interior neurons, there is no connectivity remodeling; the connection density between interior neurons is *pi_n_ţ* = 1 and the weights are slightly smaller.

### Binary model

The dynamics of the binary model are given by

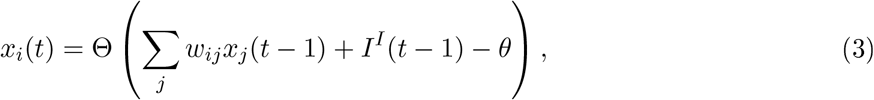

where *x_i_*(*t*) ∈ {0, 1}, is the activity of the excitatory neuron *i, i* ∈ {1,…,*N*}, at time step *t, w_ij_* the weight matrix of excitatory-to-excitatory connections, *θ* a neuron firing threshold and Θ the Heaviside step function. If present, periphery neurons are not interconnected. There are no self-connections. *I*^I^ (*t* ‒ 1) is the inhibitory input. We use global inhibition, which averages over the feedback inhibition generated by each excitatory spike, *I*^1^ (*t*) = –Θ (1/*N* ∑_j_ *x_j_*(*t*) – *θ*). At every time step each neuron is stimulated with probability *p*_sp_ to spike spontaneously. The Hebbian learning rule, applied in every step, has the form Δ*w_ij_*(*t*) = *η*(*w_ij_*)(*x_i_*(*t*) – *μ_i_*)(*x_j_*(*t*) – *μ_j_*), with a long-term average *μ_i_* of the activity of neuron *i*, and learning rate *η*(*w_ij_*). *η_j_*(*w_ij_*) = *η*_weak_ if *w_ij_* < wth and *η*_strong_ otherwise. The values of *η*_weak_ and wth differ between interior and periphery neurons, Supplementary Fig. 1b. All weights *w_ij_* are bounded between 0 and the same *w*_max_. After updating the weights with the learning rule, first the outputs and then the inputs of all neurons are normalized to *w*_sum_ = 1.

### Random walk model

The switching dynamics of a neuron in a binary network with two assemblies can be described by a one-dimensional effective random walk model. For this, we ignore the internal structure of assemblies, all details about spike mechanisms and the effect of the switching neuron on assembly activity. This reduces the spiking dynamics to randomly occurring reactivations of the assemblies and evoked or spontaneous spiking of the neuron. The weight dynamics reduce to those of the total synaptic weight *w*_1_ that the considered neuron receives from assembly 1; the input from assembly 2 is *w*_2_ = 1 – *w*_1_, due to homeostatic normalization. For simplicity, we take into account only those changes to *w*_1_ that occur when the neuron spikes together with reactivation of one of the assemblies; i.e. we neglect the small effects of weight changes between these events and of simultaneous spiking of the neuron with both assemblies. The dynamics of *w*_1_ thus become a discrete-time random walk process with statedependent noise strength, where time steps with changes in *w*_1_ correspond to the spiking of one of the assemblies together with the neuron. We use time steps of 30 ms, twice the length of those of the binary model, since reactivating assemblies are usually highly active for two consecutive time steps, Fig. 8c. We adjust spiking probabilities accordingly. Each assembly is spontaneously active in a time step with probability *p*_asbly_. If assembly 1 reactivates and *w*_1_ is larger than the spike threshold θ, the neuron also becomes active. Assembly 2 has the same effect if *w*_2_ > *θ*. Further, the neuron can be spontaneously active; this happens with probability *p*_sp_ and enables the transitions between assemblies. If the neuron and assembly 1 or assembly 2 are active together, the weight from the active assembly is potentiated, by the amount *P*(*w*_1_) or *P*(*w*_2_) prior to normalization. The actual change in the weight *w*_1_ including the effect of divisive normalization is

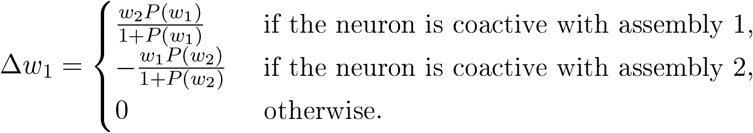

We set *P*(*w*) = *P*_weak_, if *w* < *w*_th_ and *P*(*w*) = *P*_strong_ otherwise, analogously to the binary model. The random walk model has six parameters. The neuron spontaneous spike probability *p*_sp_, the spike threshold *θ*, the threshold wth, and the ratio *P*_weak_/*P*_strong_ are set to values matching the binary model. The probability of an assembly spike *p*_asbly_ is set to that observed in simulations of the binary model. Finally, the plasticity magnitude *P*_strong_ is chosen such that the switching dynamics resemble those of the binary model.

### Statistics of weight changes

To elucidate the mechanisms underlying switching, in Fig. 5 we measure average weight changes between neurons and assemblies and their fluctuations. We record for an interior neuron the change Δ*w*_1_ of its total input *w*_1_ from assembly 1, in successive time intervals, which equal the average single neuron interspike interval. We repeat the process for all interior neurons and their inputs from all assemblies. The weights are normalized by *w*_sum_, such that *w*_1_ is between 0 and 1. We bin *w*_1_ into 50 bins of size 0.02 and calculate for each bin the average 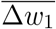 and standard deviation Std(Δ*w*_1_) of the changes ensuing those *w*_1_ that fall in it. To obtain the potential *U*(*w*_1_), we think of the average 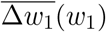 as being evoked by a force, 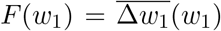, like in a classical mechanics system where friction dominates over negligible inertia. The potential *U*(*w*_1_) determines the force by *F*(*w*_1_) = −*dU*(*w*_1_)/*dw*_1_ (gradient system). *U*(*w*_1_) is computed by integrating 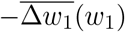 over *w*_1_.

### Model for fear memory representation

The model of fear memory representations is implemented with 150 excitatory binary neurons. The 117 interior neurons initially realize three assemblies of equal size. Each assembly has 11 periphery neurons. One assembly is chosen to represent the fear memory. Its periphery neurons are split into two context input neurons, six tone input neurons and three output neurons. At fear conditioning and each of the first recalls 100 samples of 4 interior neurons of the current fear memory realization are selected. These will be activated by the photostimulation during testing. The final recall is modeled as detection of the fear memory assembly on day 28. Thereafter the overlaps with previous assembly realizations are computed (cf. Fig. 6b) and plasticity is turned off to test the response of the system under different conditions (Fig. 6c,d). Each test lasts for 5 time steps (75ms), during the first three steps the appropriate input and photostimulated neurons are active. The circuit is considered to produce a relevant output signal, if all output neurons spike together in at least one of the last four time steps of the test. For each experimental condition, we compute the probability of such an output by averaging over 50 testing runs, for each sample of photostimulated neurons and these samples. We simulate and evaluate overall five different realizations of the system to emulate experiments with five different animals. Fig. 6 shows the means of overlaps and output activation probabilities taken over these realizations, together with their standard errors.

### XOR gate

The XOR gate model is setup as follows: The gate inputs and outputs are represented by groups of periphery neurons. The hidden layer consists of excitatory interior neurons and an inhibitory population. The interior neurons form two drifting assemblies. One of them connects to the XOR gate periphery neurons, the other is unrelated to the computation. The inhibitory population receives input from the periphery and the interior neurons; it inhibits all neurons in the network. Specifically, our system consists of 100 binary neurons, split into two assemblies of 36 neurons with 14 periphery neurons each. All inhibitory populations are modeled by single units. The XOR assembly has two input ensembles, each with 6 neurons, and 2 output neurons. Two additional inhibitory populations are added to the XOR input neurons. These receive excitation from one of the two input ensembles and get activated if all their inputs spike. If they are active, they inhibit both input ensembles. This additional inhibition is introduced to prevent the the activation of the inputs by the assembly, which would lead to both inputs being activated when the output is activated. Time steps are 15ms long.

### Linear Poisson model

Linear Poisson neurons provide good models for responses and correlations during background activity. This is because each spike has the same impact, independent of the current state of the neuron. This fits to background activity, where the state of a neuron stays close to a baseline all the time. Linear Poisson neurons are stochastically spiking neurons with instantaneous rates *f_i_*(*t*), *i* = 1,…, *N*, evolving in continuous time. The rates are excited by spikes from the other neurons in the network and follow the linear dynamics

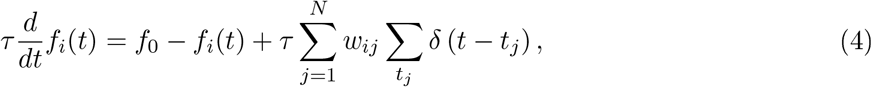

where *f*_0_ is a constant spontaneous rate due to the assumed embedding in a fluctuation-driven asynchronous irregular activity state. A spike from neuron *j* increases *f_i_*(*t*) in a jump-like manner by a nonnegative synaptic weight *w_ij_*. Between input spikes *f_i_*(*t*) decays exponentially with time constant *τ* = 10ms to the spontaneous rate *f*_0_ = 0.75Hz. On average a spike of neuron j induces *τw_ij_* additional spikes in neuron *i*. Global or explicitly modeled inhibition is assumed to be implicitly contained in the model; it contributes to the randomness of spike generation, both of spontaneous and of excited activity. The network dynamics can be solved in an event-based fashion allowing for very long simulation times.

The synapses in our Poisson networks change according to the following plasticity rule. When a neuron spikes, its existing input and output synapses are changed depending on the current level of excitation of the corresponding partner neurons, which we measure by the instantaneous rate above baseline, *f_i_*(*t*) – *f*_0_ The dependence is given by a function Δ*w*(*f_i_* – *f*_0_), which is negative (giving rise to LTD) for small and average values of the excitation and positive (giving rise to LTP) for larger values; for simplicity we use a quadratic function (see Supplementary Fig. 1) with Δ*w*(0) = 0. The weights are bounded by 0 and *w*_max_. We model homeostatic plasticity by input and output normalization of summed synaptic weights as in the other network models. Here this implies that the average number of additional spikes in the network induced by a spike of neuron *j* (also referred to as ‘branching parameter’) remains constant: *τ*∑_*i*_ *w_ij_* = *τw*_sum_ = 0.25. The synaptic connections in our Poisson networks also remodel spontaneously as described above.

### Correlations and reactivation amplitudes

To quantify the pairwise spike correlations in our network models in Fig. 8 first and second column, we use the Pearson correlation coefficients of spike counts [41] in time bins of size 150ms. We measure the spike counts in simulations of static networks with connectivity and weights taken from the plastic simulations exhibiting drifting assemblies after the first complete remodeling. The measurement time is 2 h for the LIF, 10 h for the Poisson and 15 h for the binary network. The neurons of the static networks are sorted according to their detected assembly membership as in the figures demonstrating drifting assemblies. This allows us to partition the neuron pairs into intra-assembly pairs (where both neurons are in the same assembly) and inter-assembly pairs (where each neuron is in a different assembly) and to show their contributions to the distributions of correlation coefficients in the networks (Fig. 8, second column).

To investigate spontaneous assembly reactivation in Fig. 8 third column we consider the summed spiking activity of all neurons in an assembly. This assembly spiking activity is then temporally filtered with a moving time window of size 50 ms for the LIF, 75 ms for the binary and 100 ms for the Poisson model. Reactivation events in our LIF and binary networks are shorter than the window size. Further, the analytical duration distribution [38] shows that for our Poisson networks’ branching parameter and time constant only 0.013% of single avalanches are longer than 100 ms, justifying the latter window size. The filtering results in a time series, which gives at each time the number of spikes that have occurred in an assembly during the preceding time window. We locate local maxima of this time series to detect putative assembly reactivation. If two local maxima are found with a temporal distance less than the filter window size, only the larger one is kept. The heights of the local maxima are initially numbers of spikes; we normalize them by the corresponding assembly size and call them (relative) reactivation amplitudes. We only consider amplitudes above or equal to a minimal value (0.2 for LIF, 0.15 for binary and 0.25 for Poisson networks). Fig. 8 third column shows histograms of the amplitudes obtained from the same simulations as the correlation measurements. Dividing the counts of different amplitudes by the measurement time gives the displayed occurrence rate. The associated complementary cumulative distributions (see Fig. 8, third column insets) indicate the rates at which putative assembly reactivation with sizes exceeding the given amplitude occur.

## Acknowledgements

We thank Paul Züge for fruitful discussions, Hans Günter Memmesheimer for help with the graphical illustration and Jonas Nitzsche for providing the photo of the thread in Fig. 1, and the German Federal Ministry of Education and Research (BMBF) for support via the Bernstein Network (Bernstein Award 2014, 01GQ1710).

## Notes

### Competing Interest Statement

The authors have declared no competing interest.

