## Supplementary Figures and Notes for "Drifting Assemblies for Persistent Memory"

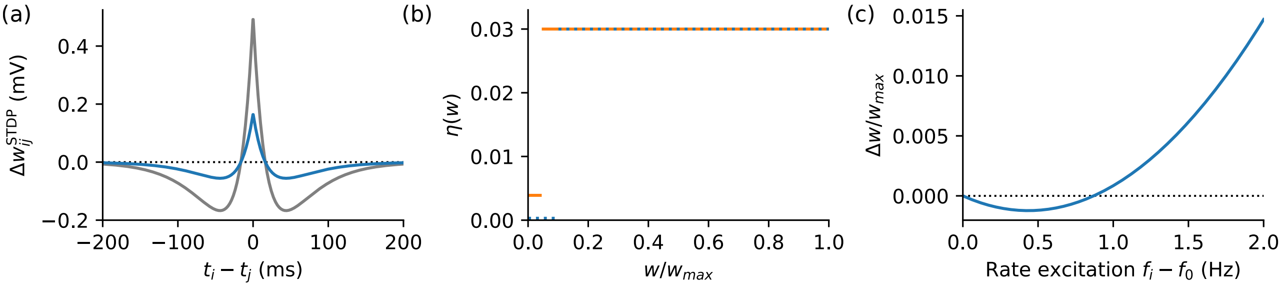


*Figure S1: Plasticity rules for different models. (a) STDP window used in the LIF model with (blue, main text Fig. 2), and without (gray, main text Fig. 3) spontaneous connectivity remodeling. The weight change is given in terms of the change of the peak EPSP that a synapse evokes in a resting neuron. (b) Weight dependence of the learning rate in the binary model (Fig. S4), for interior (dashed blue) and for periphery (solid orange) neurons. (c) Dependence of the weight update on the excitation level in the Poisson model (Fig. S6). Black dotted lines in (a) and (c) indicate border between depression and potentiation.*

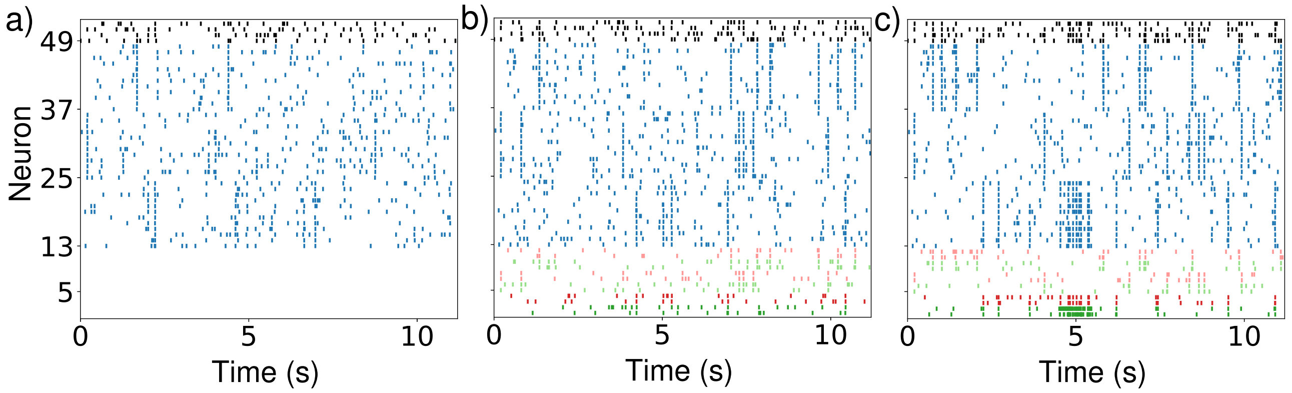


*Figure S2: Associative memory property and basic input-output functionality of assemblies. (a) Assemblies spontaneously activate due to sparse background activity and in absence of periphery neuron spiking. (b) In presence of periphery neurons, the rate of activation is higher. (c) Stimulation of a pair of input neurons (green, neurons 1,2) stimulates their assembly (assembly 1), which specifically stimulates its output neurons (red, neurons 3,4), demonstrating basic functionality of our circuit. The spike trains are sorted according to the assemblies that the neurons belong to at* $t = 0s$. *The input neurons to assemblies 1,2 and 3 have indices 1,2 (green), 5,6 (light green) and 9,10 (light green). The output neurons of assemblies 1,2 and 3 have indices 3,4 (red), 7,8 (light red) and 11,12 (light red). The first twelve assembly neurons of each assembly are displayed in blue, with indices 13-24, 25-36 and 37-48. Further, the spike trains of four inhibitory neurons are shown in black.*

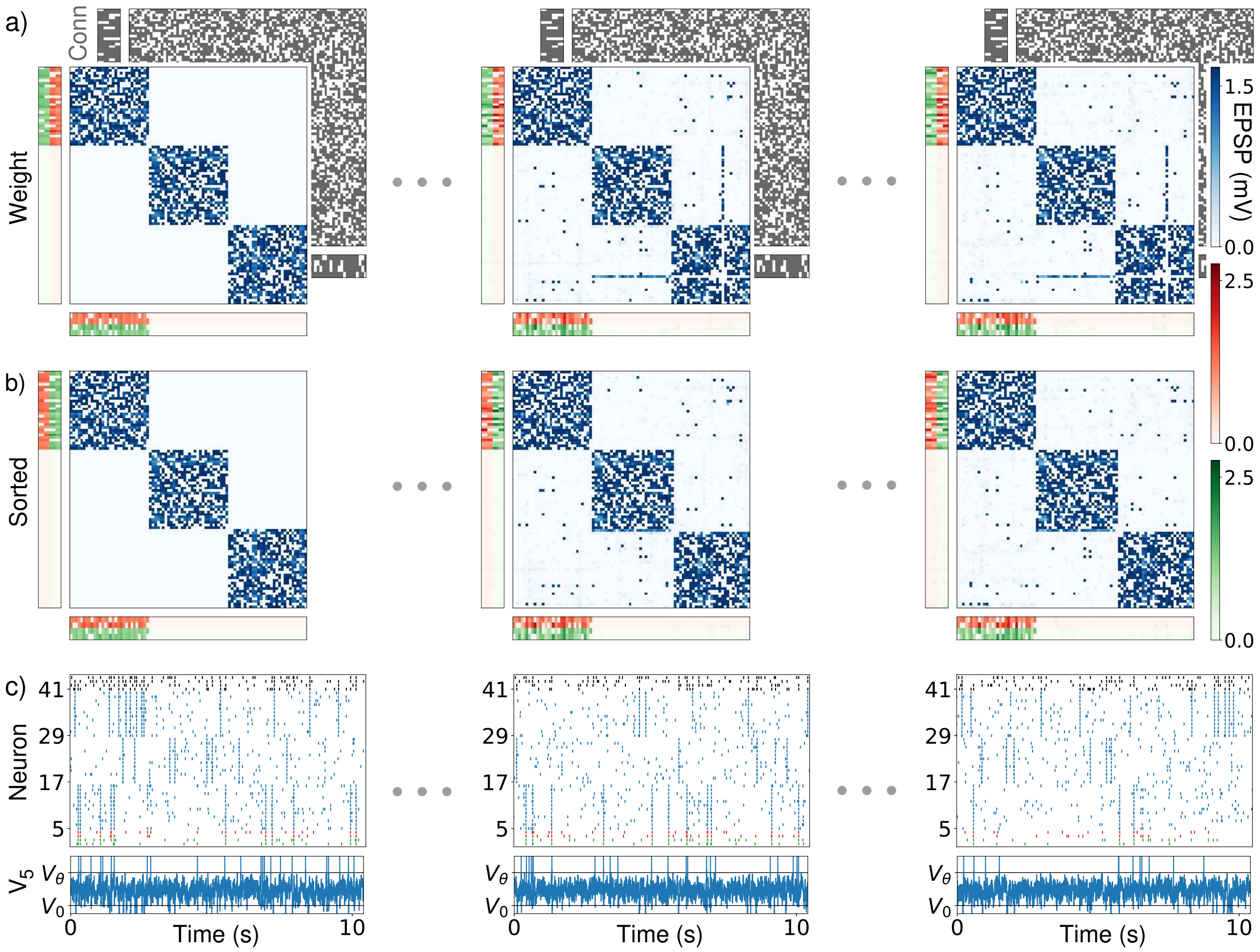


*Figure S3: LIF network as in main text Fig. 2 but without spontaneous remodeling. Panels (a-c): like panels (b-d) in Fig. 2, but for 0, 2 and 99 hours of simulated time. The connectivity matrix is constant. After an initial phase of partial remodeling (compare the first and second column), no more neurons change assembly (compare the second and third column).*

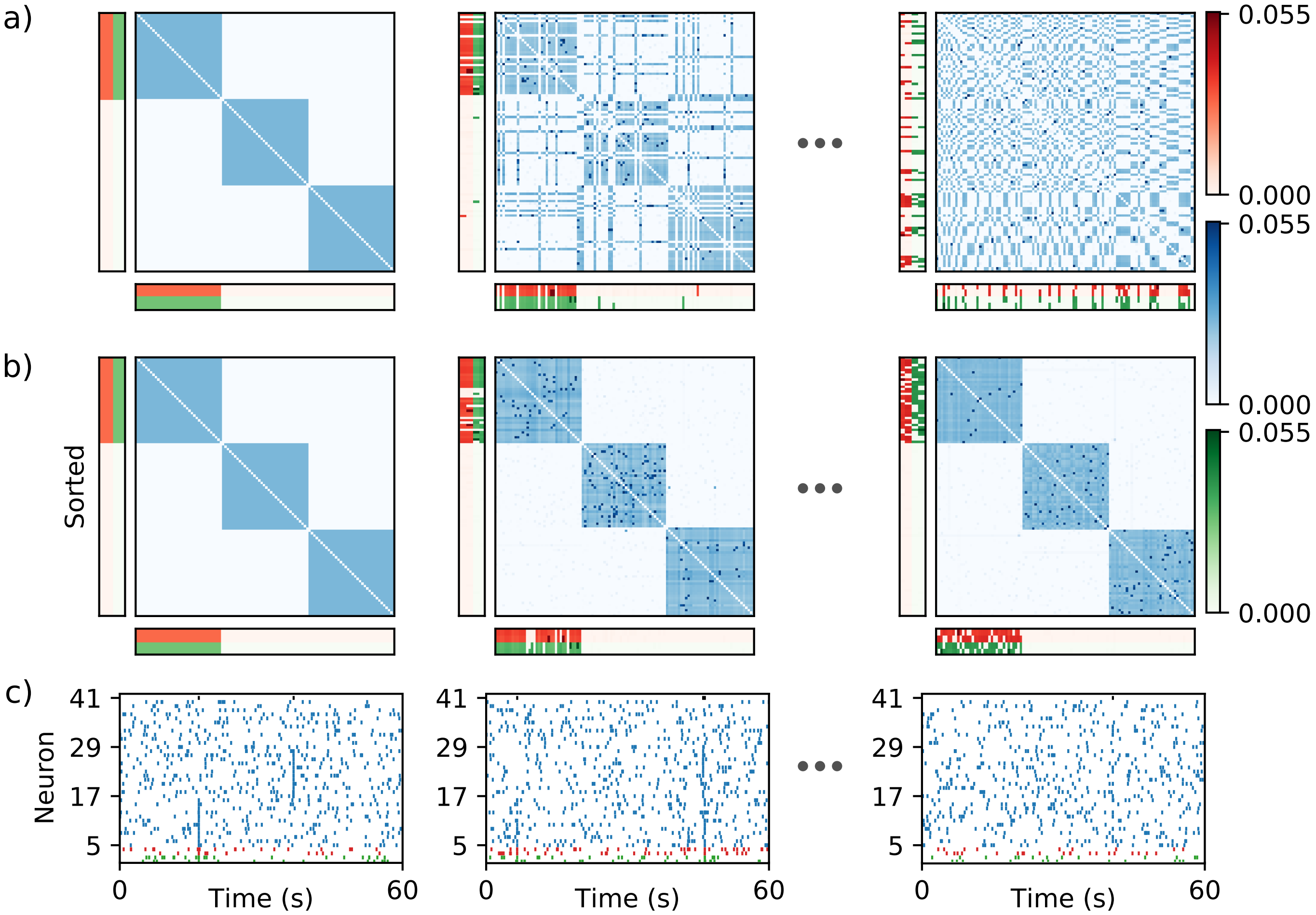


*Figure S4: Drifting assemblies in a network of binary neurons without spontaneous remodeling. Display is like in main text Fig. 4, but with periphery neuron weights as in main text Fig. 2. First column: Network initialization with three assemblies. Second column, after one day: several interior neurons switched to a new assembly. Third column, after 18 days: The assemblies have drifted away, the weight matrix has completely remodeled. (d) shows spike trains of the input (green) and output (red) neurons of assembly 1, of 12 neurons from each of the ensembles that initially form assembly 1 (5-16), 2 (17-28) and 3 (29-40) and of the inhibitory unit (black).*

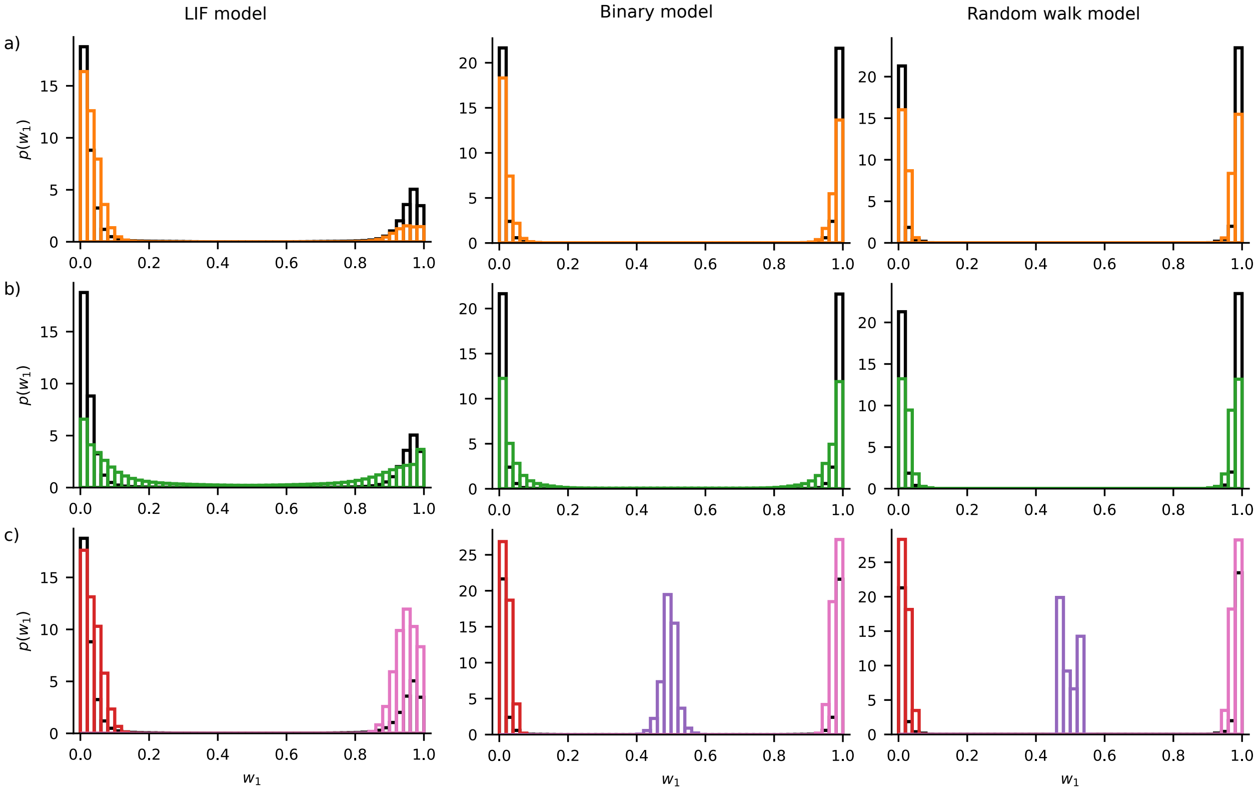


*Figure S5: Stationary distributions of $w_1$ for the LIF network (left, black), the binary network (middle, black), and the random walk (right, black) models, compared to the occupancy distributions of related Markov simulations. (a) Markov simulation accounting for drift and noise (orange) (b) Markov simulation accounting for noise only (green) (c) Markov simulation with drift and homogenized noise. For homogenized noise, the distribution of the Markov simulations depend on the initial $w_1$, since switching does not occur within the used simulation time and the neuron stays within one of the two (LIF) or three (binary, random walk) potential valleys. For the Markov simulation to the LIF model the initial values are $w_1(0) = 0$ (red) and $w_1(0) = 1$ (pink). For the binary and the random walk model they are $w_1(0) = 0$ (red), $w_1(0) = 1$ (pink), and $w_1(0) = 0.5$ (purple).*

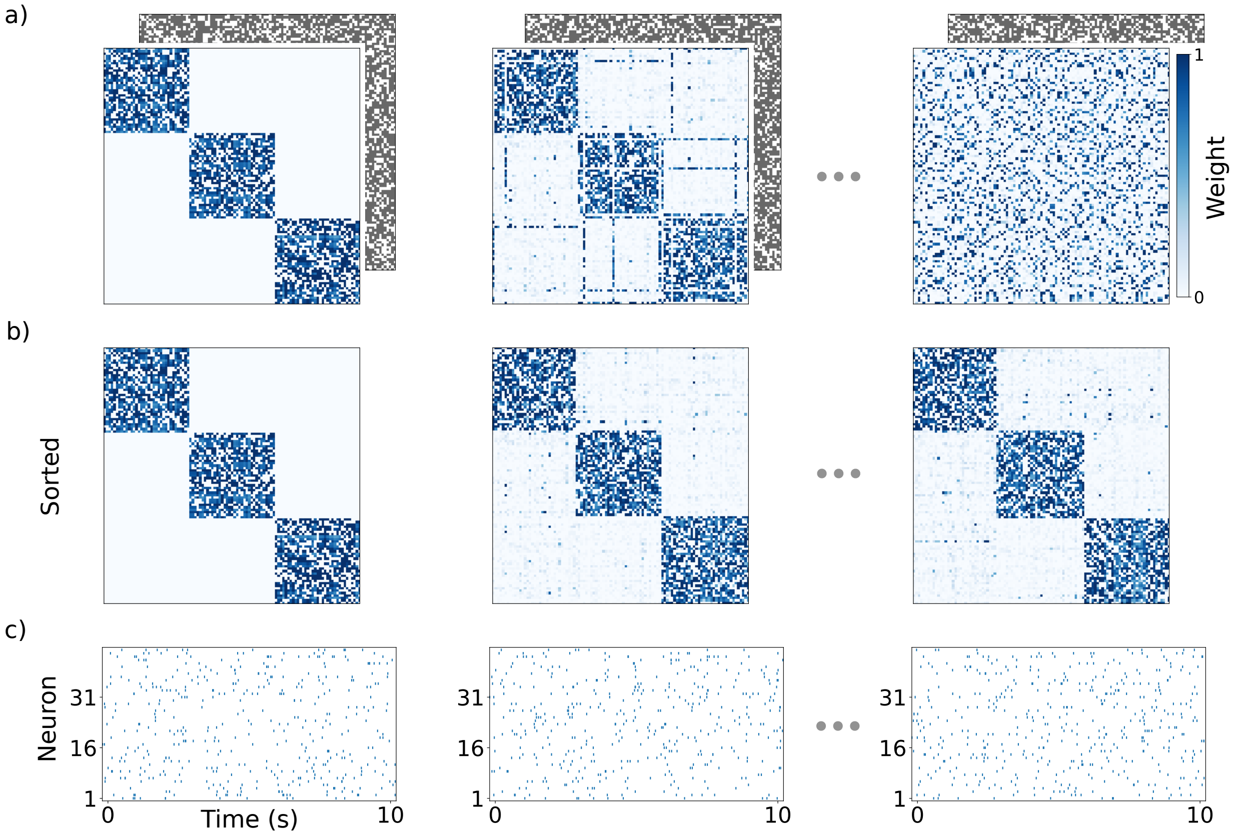


*Figure S6: Drifting assemblies in a network of linear Poisson spiking neurons with spontaneously remodeling connectivity and without periphery neurons. Panels (a-c): like panels (b-d) in Fig. 2, but for 0, 72 and 1440 hours of simulated time, i.e. the second column shows the network after three days and the third column after 60 days. The underlying connectivity matrix is spontaneously remodeling and drives the drift. The weights are normalized by their maximal possible value* $w_{\max}$. *(c) Spike trains of 15 neurons from each of the ensembles that initially form assembly 1 (1-15), 2 (16-30) and 3 (31-45). Spiking activity is asynchronous and irregular without visible assembly reactivation.*

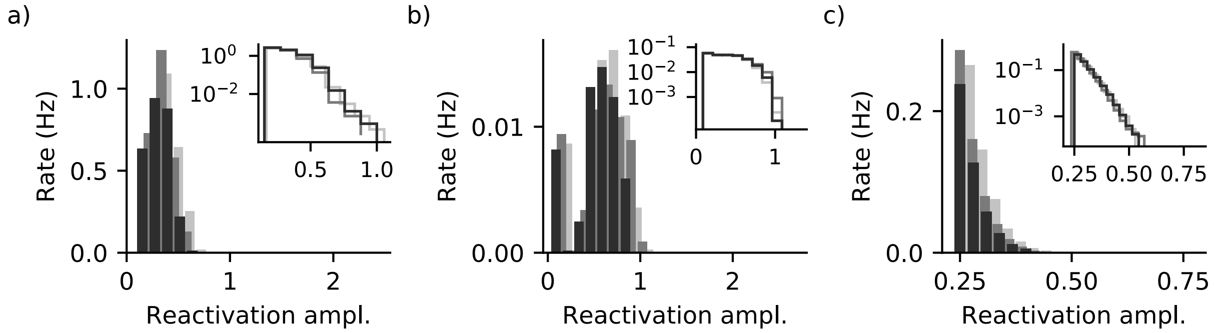


*Figure S7: Distributions of reactivation amplitudes in our network models as in main text Fig. 8, third column, but for randomly selected neuron ensembles of the same size as the three assemblies in the networks. For the LIF (a) and the binary (b) model the histograms consist of amplitudes of about* $1/3$ *of the typical assembly reactivation amplitudes visible in main text Fig. 8, third column. This fraction agrees with the expected neuron overlap of the random ensembles with the assemblies. The observation therefore confirms the confinement of reactivations to single assemblies, which is also apparent from the sorted spiking dynamics (cf. the first columns of Figs. 2, 4, S3, S4 and Fig. S2). For the linear Poisson network model (c) the amplitude distribution appears to exponentially decay in a similar fashion as for the assemblies in the network (main text Fig. 8c right). The semi-logarithmic plot of the complementary cumulative distribution in the inset reveals that for randomly selected ensembles the exponential decay is slightly faster.*

### Supplementary Notes

##### Family resemblance and identity

L. Wittgenstein uses the analogy of a thread to clarify the relation of the different objects denoted by a word. They need not be directly related by overarching commonalities, but often possess many partial, overlapping “family resemblances”, as explained in paragraphs 66 and 67 of ref. [1]:

“66. Consider, for example, the activities that we call “games”. I mean board-games, card-games, ball-games, athletic games, and so on. What is common to them all? – Don’t say: “They must have something in common, or they would not be called ‘games’ ” – but look and see whether there is anything common to all. – For if you look at them, you won’t see something that is common to all, but similarities, affinities, and a whole series of them at that. (...)

67. I can think of no better expression to characterize these similarities than “family resemblances”; for the various resemblances between members of a family – build, features, colour of eyes, gait, temperament, and so on and so forth – overlap and criss-cross in the same way. – And I shall say: ‘games’ form a family. And likewise the kinds of number, for example, form a family. Why do we call something a “number”? Well, perhaps because it has a – direct – affinity with several things that have hitherto been called “number”; and this can be said to give it an indirect affinity with other things that we also call “numbers”. And we extend our concept of number, as in spinning a thread we twist fibre on fibre. And the strength of the thread resides not in the fact that some one fibre runs through its whole length, but in the overlapping of many fibres.”

Different parts consisting of different ensembles of fibers are parts of the same thread. The overlaps of the fibers give rise to identity over space. Likewise in our model the temporal overlaps of neurons participating in the different assembly realizations give rise to the identity of the memory over time. The various family resemblances are in our model the similarities of neuron ensembles forming the assemblies at close-by times. We note that the thread analogy is sufficiently flexible to cover the change of memories by learning new facts, for example about apples: If a thread becomes larger, of a different material or loosely intertwined with other threads, it still maintains its identity.

The question of identity was already a central one in ancient Greek philosophy. Heraclitus of Ephesus saw objects in continuous change over time. Well known are aphorisms that shall paraphrase his ideas such as “It is not possible to step twice into the same river” [2, 3], with a continuation: “or to come into contact twice with a mortal being in the same state”. Another famous aphorism of this kind is cited at the end of our main text.

The ancient historian Plutarch recounts a puzzle on identity, the “Ship of Theseus”: After its famous trip this ship is displayed in Athens. Over time one part after the other is gradually replaced. At some point no piece is original anymore. Is it nevertheless Theseus’ ship? T. Hobbes added a twist, suggesting that the original parts might be collected somewhere upon their replacement. If in the end they are put together again to a ship, which one is now Theseus’ ship [4]? This is another analogy to the assembly model that we propose. The question of identity is for our assemblies solved by the input and output neurons: They connect to the neuron ensemble that forms the memory representation. Suppose that after some time all neurons that originally formed, for example, the apple assembly have been exchanged. Theis assembly now consists of different neurons (like the displayed ship of different parts), but it is the apple assembly, since the right input and output neurons connect to it; the new neuron ensemble forming the assembly mediates the correct behavior. Now assume that the neurons that originally formed the apple assembly connect later, for some reason, to an assembly again (the original ship parts are put together again). This assembly will not be the apple assembly, since suitable input and output neurons are not connected to it.

#### Associative memory property and input-output functionality of assemblies

This section checks that assemblies of LIF neurons have the associative memory property and that their circuits of inputs, assemblies and outputs have basic input-output functionality. For the former, we simulate the dynamics in absence of periphery neuron activity, Fig. S2a. We observe spontaneous assembly reactivation on a background of sparse activity. Since the sparse background activity will only activate part of an assembly, this shows that partial activation can lead to recall (complete or near complete assembly reactivation) as required for associative memory. In presence of periphery neurons, spontaneous assembly reactivation is more frequent, Fig. S2b. This is expected because the periphery neurons contribute spontaneous spikes and help amplifying assembly activity. Further our circuits are functional in the sense that sufficiently strong stimulation of input neurons activates an assembly, which in turn activates its output neurons, Fig. S2c. Activation is specific, other assemblies and periphery neurons do not increase their activity. All simulations in Fig. S2 are done after the first complete restructuring of the system was detected. We set this time to $t = 0$s. In Fig. S2a,b the coupling and connectivity matrices are kept constant at their values at $t = 0$s. This avoids compensation of the missing periphery neurons in Fig. S2a. Fig. S2b shows the beginning of the simulation analyzed in main text Fig. 8a. In Fig. S2c external stimulation forces the input neurons to generate 50Hz Poisson spike trains for 1s after $t = 4.5$s. The resulting spiking activity does not destroy the circuit structure.

#### Static assemblies in absence of spontaneous synaptic remodeling

If the synaptic connectivity of the network in main text Fig. 2 is kept constant, the assemblies do not drift, see Fig. S3. There is an initial phase where the assemblies adapt to the connectivity matrix: neurons that receive few input connections from their assembly leave it. In the simulation of Fig. S3, one neuron changes assembly at around one hour simulated time, see also main text Fig. 3. No more changes occur until the end of the simulation at 100 hours.

#### Markov simulation of neuron transitions between assemblies

We use the sampled mean $\overline{\Delta w_1}(w_1)$ and standard deviation $\mathrm{Std}(\Delta w_1)(w_1)$ of weight changes to compare the contributions of drift and noise to neuron transitions between assemblies for different models. Since the sampling interval is comparably large, we assume that the change in $w_1$ depends only on its previous value (Markov assumption). For simplicity we further assume that noise is normally distributed. We therefore simulate the weight dynamics as $w_1(t+1) = \overline{\Delta w_1}(w_1(t)) + \xi(w_1(t))$, where $\xi(w_1(t)) \sim \mathcal{N}(0, \mathrm{Std}(\Delta w_1)(w_1(t)))$, and $w_1$ is clipped to the interval $[0,1]$ after each step. We find that the stationary probability density functions of $w_1$ for the full models, $p_{\mathrm{full}}(w_1)$, and those of the corresponding Markov simulations are in acceptable agreement, Fig. S5a. Deviations are likely due to the non-Gaussianity of the weight change distributions in the network simulations. To study the contribution of noise, we repeat the Markov simulations, but without including the effect of the mean, $w_1(t+1) = \xi(w_1(t))$. We find that this process also agrees well with the full models, Fig. S5b. Finally, to examine the contribution of mean, we calculate the average noise standard deviation, $\overline{\mathrm{Std}(\Delta w_1)(w_1)} = \int_0^1 \mathrm{Std}(\Delta w_1)(w_1) p_{\mathrm{full}}(w_1) dw_1$, and simulate the dynamics with state-independent noise $\xi \sim \mathcal{N}(0, \overline{\mathrm{Std}(\Delta w_1)(w_1(t))})$. With this noise, the neuron does not leave its potential valleys (cf. main text Fig. 5c) within the simulated periods. For the LIF model, the neuron thus stays with the assemblies, for the binary model it stays with the assemblies or gets trapped in the middle of transition. Thus, also for homogenized noise the dynamics show crucial aspects of the full dynamics. We conclude that both drift and noise inhomogeneity contribute to the switching dynamics. The results further indicate that the inhomogeneous noise is already sufficient to generate the crucial features of the stationary distribution, the meta-stable states and the switching. Noise-induced multistability has been observed in different models and natural systems before, e.g. for electrical and chemical oscillations, populations dynamics and foraging behavior [5, 6, 7, 8].

#### Parameters of models used for the simulations

##### LIF model with connectivity remodeling

*Neuron numbers*: excitatory neurons: $N_E = 102$; interior neurons: $N_{\text{int}} = 90$; periphery neurons: 12; inhibitory neurons: $N_I = 20$.

*Network structure*: connection probability between interior neurons: $p_{\text{int}} = 0.6$; connection probability between interior and periphery neurons: $p_{\text{peri}} = 0.8$; periphery neurons have no connections between each other; connection probability between excitatory and inhibitory neurons and between inhibitory and inhibitory neurons: 1; there are no self-connections of neurons.

*Neuron parameters*: spike threshold: $V_\theta = 20\text{mV}$; reset potential $V_0 = 0\text{mV}$; resting potential $V_{\text{rest}} = 10\text{mV}$; membrane time constant: $\tau_m = 10\text{ms}$; absolute refractory period: $\tau_{\text{ref}} = 5\text{ms}$; sum of input and sum of output weights of an interior neuron: $w_{\text{sum}} = 253.125\text{mV} = \frac{3}{4}\left[(N_{\text{asbly}} - 1)\, p_{\text{int}} w_{\text{max}} + N_{\text{peri}} p_{\text{peri}} w_{\text{max,peri}}\right]$, the angular bracketed term is the expected input of an interior neuron from a typical size assembly and its periphery neurons, if all weights were at their individual maximum; sum of input and sum of output weights of a periphery neuron: $w_{\text{sum,peri}} = 225.0\text{mV} = \frac{1}{4}\left[N_{\text{asbly}} p_{\text{peri}} w_{\text{max,peri}}\right]$, the angular bracketed term is the expected input of a periphery neuron from a typical size assembly, if all weights are at their individual maximum; noise input strength: $\sigma = 3.5\text{mV}$.

*Excitatory synapses*: time constant: $\tau_E = 2\text{ms}$; maximal synaptic strength of synapses between interior neurons: $w_{\text{max}} = 12.5\text{mV}$, evoking a peak EPSP of 1.67mV in a resting postsynaptic neuron; maximal synaptic strength of synapses between interior and periphery neurons: $w_{\text{max,peri}} = 37.5\text{mV}$, evoking a peak EPSP of 5.02mV in a resting postsynaptic neuron; strength of synapses to inhibitory neurons: $w_{\text{E}\to\text{I}} = 4.96\text{mV}$, evoking a peak EPSP of 0.66mV in a resting postsynaptic neuron.

*Inhibitory synapses*: time constant: $\tau_I = 5\text{ms}$; strength of synapses to excitatory neurons: $w_{\text{I}\to\text{E}} = -5.06\text{mV}$, evoking a peak inhibitory postsynaptic potential (IPSP) of $-1.27\text{mV}$ in a resting postsynaptic neuron; strength of synapses to inhibitory neurons: $w_{\text{I}\to\text{I}} = -5.33\text{mV}$ evoking a peak IPSP of $-1.33\text{mV}$ in a resting postynaptic neuron.

*STDP window*: $\Delta w_{ij}(\Delta t) = \frac{\eta}{a - b(1+\delta)}\left[a \exp(-a\,|\Delta t|) - b(1+\delta)\exp(-b\,|\Delta t|)\right]$, where $\Delta t = t_i - t_j$ is the time difference between the postsynaptic and the presynaptic spike; window amplitude: $\eta = 1.25\text{mV}$, LTP peak at 0ms; LTP decay rate: $a = \frac{1}{\tau_{\text{LTP}}} = \frac{1}{20\text{ms}}$; LTD decay rate: $b = \frac{1}{\tau_{\text{LTD}}} = \frac{1}{40\text{ms}}$; ratio of integrated LTD and LTP: $1 + \delta = 1 + \frac{1}{3}$.

*Connectivity change*: life and absence time of synapses between interior neurons: $L_{\text{int}} = 2000\text{s}$ and $A_{\text{int}} = 1333.3\text{s}$; life and absence time of synapses between interior and periphery neurons: $L_{\text{peri}} = 2000\text{s}$ and $A_{\text{peri}} = 500\text{s}$.

*Memory representation*: number of assemblies: 3; initial number of interior neurons per assembly: $N_{\text{asbly}}(0) = 30$; periphery neurons per assembly: $N_{\text{peri}} = 4$.

*Simulation*: time step: 0.25ms; total simulated time: 100 hours.

##### LIF model without connectivity remodeling

Same parameters as for simulations with connectivity remodeling with the following exceptions:

*Neuron numbers*: excitatory neurons: $N_E = 102$; interior neurons: $N_{\text{int}} = 102$; periphery neurons: 0; inhibitory neurons: $N_I = 20$.

*Network structure*: connection probability between interior neurons: $p_{\text{int}} = 1$.

*Neuron parameters*: sum of input and sum of output weights of an interior neuron: $w_{\text{sum}} = 247.5\text{mV} = \frac{3}{5}\left[(N_{\text{asbly}} - 1)\, p_{\text{int}} w_{\text{max}}\right]$.

*Excitatory synapses*: strength of synapses to inhibitory neurons: $w_{\text{E}\to\text{I}} = 4.85\text{mV}$, evoking a peak EPSP of 0.65mV in a resting postsynaptic neuron.

*Inhibitory synapses*: time constant: strength of synapses to excitatory neurons: $w_{\text{I}\to\text{E}} = -4.95\text{mV}$, evoking a peak inhibitory postsynaptic potential (IPSP) of $-1.24$mV in a resting postsynaptic neuron; strength of synapses to inhibitory neurons: $w_{\text{I}\to\text{I}} = -5.21\text{mV}$ evoking a peak IPSP of $-1.30$mV in a resting postynaptic neuron.

*STDP window*: window amplitude: $\eta = 3.75\text{mV}$.

*Connectivity change*: No connectivity change.

*Memory representation*: initial number of interior neurons per assembly: $N_{\text{asbly}}(0) = 34$.

*Simulation*: total simulated time: 50 hours.

##### Binary model

*Neuron numbers*: excitatory neurons: $N_E = 120$; interior neurons: $N_{\text{int}} = 108$; periphery neurons: 12.

*Network structure*: connection probability between all neurons: $p = 1$; there are no self-connections of neurons.

*Neuron parameters*: spike threshold: $\theta = 0.1$; spontaneous spike probability $p_{\text{sp}} = 0.004$; sum of input and sum of output weights of a neuron: $w_{\text{sum}} = 1$; maximal synaptic strength $w_{\text{max}} = 0.0556$.

*Learning Rule*: learning rate for strong synapses $\eta_{\text{strong}} = 0.03$; learning rate for weak synapses between interior neurons $\eta_{\text{weak,int}} = 0.0039$; learning rate for weak synapses between interior and periphery neurons $\eta_{\text{weak,peri}} = 0.0009$; weak synapse cutoff for synapses between interior neurons $w_{\text{th,int}} = 0.05 w_{\text{max}}$; weak synapse cutoff for synapses between interior and periphery neurons $w_{\text{th,peri}} = 0.1 w_{\text{max}}$.

*Connectivity change*: No connectivity change.

*Memory representation*: number of assemblies: 3; initial number of interior neurons per assembly: $N_{\text{asbly}}(0) = 36$; periphery neurons per assembly: $N_{\text{peri}} = 4$.

*Simulation*: time step: 15ms; total simulated time: 86 days.

##### Binary model for transition mechanism analysis

*Neuron numbers*: excitatory neurons: $N_E = 72$; interior neurons: $N_{\text{int}} = 72$; periphery neurons: 0.

*Network structure*: connection probability between all neurons: $p = 1$; there are no self-connections of neurons.

*Neuron parameters*: spike threshold: $\theta = 0.1$; spontaneous spike probability $p_{\text{sp}} = 0.004$; sum of input and sum of output weights of a neuron: $w_{\text{sum}} = 1$; maximal synaptic strength $w_{\text{max}} = 0.0556$.

*Learning Rule*: learning rate for strong synapses $\eta_{\text{strong}} = 0.03$; learning rate for weak synapses between interior neurons $\eta_{\text{weak,int}} = 0.0045$; weak synapse cutoff for synapses between interior neurons $w_{\text{th,int}} = 0.05 w_{\text{max}}$.

*Connectivity change*: No connectivity change.

*Memory representation*: number of assemblies: 2; initial number of interior neurons per assembly: $N_{\text{asbly}}(0) = 36$.

*Simulation*: time step: 15ms; total simulated time: 15 days.

**Binary model of the fear memory representation**

*Neuron numbers*: excitatory neurons: $N_E = 150$; interior neurons: $N_{\text{int}} = 117$; periphery neurons: 33.

*Network structure*: connection probability between all neurons: $p = 1$; there are no self-connections of neurons.

*Neuron parameters*: spike threshold: $\theta = 0.19$; spontaneous spike probability $p_{\text{sp}} = 0.05$; sum of input and sum of output weights of a neuron: $w_{\text{sum}} = 1$; maximal synaptic strength $w_{\text{max}} = 0.04$.

*Learning Rule*: learning rate for strong synapses $\eta_{\text{strong}} = 0.01$; learning rate for weak synapses between interior neurons $\eta_{\text{weak,int}} = 0.0012$; learning rate for weak synapses between interior and periphery neurons $\eta_{\text{weak,peri}} = 0.0002$; weak synapse cutoff for synapses between interior neurons $w_{\text{th,int}} = 0.05w_{\text{max}}$; weak synapse cutoff for synapses between interior and periphery neurons $w_{\text{th,peri}} = 0.1w_{\text{max}}$.

*Connectivity change*: No connectivity change.

*Memory representation*: number of assemblies: 3; initial number of interior neurons per assembly: $N_{\text{asbly}}(0) = 39$; periphery neurons per assembly: $N_{\text{peri}} = 11$.

*Simulation*: time step: 15ms; total simulated time: 86 days.

**Binary model of the XOR gate**

*Neuron numbers*: excitatory neurons: $N_E = 100$; interior neurons: $N_{\text{int}} = 72$; periphery neurons: 28.

*Network structure*: connection probability between all neurons: $p = 1$; there are no self-connections of neurons.

*Neuron parameters*: spike threshold: $\theta = 0.1$; spontaneous spike probability $p_{\text{sp}} = 0.004$; sum of input and sum of output weights of a neuron: $w_{\text{sum}} = 1$; maximal synaptic strength $w_{\text{max}} = 0.0455$.

*Learning Rule*: learning rate for strong synapses $\eta_{\text{strong}} = 0.03$; learning rate for weak synapses between interior neurons $\eta_{\text{weak,int}} = 0.0105$; learning rate for weak synapses between interior and periphery neurons $\eta_{\text{weak,peri}} = 0.0009$; weak synapse cutoff for synapses between interior neurons $w_{\text{th,int}} = 0.05w_{\text{max}}$; weak synapse cutoff for synapses between interior and periphery neurons $w_{\text{th,peri}} = 0.15w_{\text{max}}$.

*Connectivity change*: No connectivity change.

*Memory representation*: number of assemblies: 2; initial number of interior neurons per assembly: $N_{\text{asbly}}(0) = 36$; periphery neurons per assembly: $N_{\text{peri}} = 14$.

*Simulation*: time step: 15ms; total simulated time: 86 days.

**Random walk model**

*Neuron parameters*: spontaneous spike probability $p_{\text{sp}} = 0.008$; spike threshold $\theta = 0.1$.

*Learning Rule*: plasticity magnitude $P = 0.45$; plasticity magnitude for weak synapses $P_{\text{weak}} = 0.15P$; weak synapse cutoff $w_{\text{th}} = 0.05$.

*Assembly parameters*: spike probability $p_{\text{A}} = 0.0014$.

*Simulation*: time step: 30ms; total simulated time: 347 days.

.

**Linear Poisson model**

*Neuron numbers*: $N_E = 105$; interior neurons: $N_{\text{int}} = 105$; periphery neurons: 0.

*Network structure*: connection probability between neurons: $p_{\text{int}} = 0.65$; there are no self-connections of neurons.

*Neuron parameters*: time constant: $\tau = 10$ms; spontaneous rate: $f_0 = 0.75$Hz; sum of input and sum of output weights of a neuron: $\tau w_{\text{sum}} = 0.25$, maximal synaptic weight: $\tau w_{\max} = 0.0124$.

*Plasticity rule*: symmetric weight change upon a spike of neuron $j$: $\Delta w(\Delta f_i) = 0.01/w_{\max}\,(\Delta f_i - 0.87\text{Hz})\,\Delta f_i$, where $\Delta f_i = f_i(t) - f_0$ is the current level of excitation of the post- or pre-synaptic partner neuron.

*Connectivity change*: life and absence time of synapses: $L_{\text{int}} = 6$h and $A_{\text{int}} = 3.23$h.

*Memory representation*: number of assemblies: 3; initial number of interior neurons per assembly: $N_{\text{asbly}}(0) = 35$.

*Simulation*: event-based; simulated time: 125 days.
